# ST-PARM: Pareto-Complete Inference-Time Alignment for Multi-Objective Protein Design

**DOI:** 10.64898/2026.03.17.712483

**Authors:** Rujie Yin, Yang Shen

## Abstract

**Motivation:** Protein engineering is inherently multi-objective: improving one property can degrade others, so practical workflows require generating non-dominated (Pareto-optimal) candidates spanning a trade-off surface. Linear objective scalarization and deterministic pairwise preference learning can under-explore non-convex Pareto regions and amplify noise from uncertain evaluators, limiting Pareto coverage and trade-off controllability.

**Results:** We introduce Smooth Tchebycheff Preference-Aware Reward Model (ST-PARM), an inference-time alignment framework that steers a frozen protein language model along user-specified trade-offs with a lightweight reward model trained only once. ST-PARM combines (i) a reward-calibrated pairwise preference loss that is uncertainty-aware by down-weighting ambiguous comparisons under noisy evaluators, (ii) a smooth Tchebycheff scalarization that is Pareto-complete in principle and improves empirical trade-off coverage, and (iii) latent-space pair-construction strategies. On GFP fluorescence–stability (full-length design) and IL-6 nanobody stability–solubility (CDR3+suffix design), ST-PARM delivers broader Pareto coverage and stronger preference tracking than baselines PARM and MosPro. For GFP, a conservative structural screen for local confidence and global fold preservation retains a broad frontier and strong controllability, yielding an actionable cohort for downstream assays. We also provide cross-evaluator robustness checks, a three-objective extension, and a natural-language alignment generality check in the Supplement, establishing a practical foundation for controllable sequence generation under competing multi-objectives and noisy measurements.

**Availability and Implementation:** https://github.com/Shen-Lab/ST-PARM.

**Supplementary Information:** Supplementary data are provided with the submission.

## 1. Introduction

Designing functional proteins requires balancing competing objectives—e.g., fluorescence vs. stability in reporters or affinity vs. solubility in therapeutics. Thus the goal is a *set* of non-dominated candidates along a trade-off surface (the *Pareto frontier*) rather than a single optimum [Rabia et al., 2018, Nanda et al., 2017]. Broader Pareto coverage expands the set of non-dominated candidates available to downstream selection and better accommodates shifting or unknown constraints [Hong and Kortemme, 2024].

Protein trade-offs are often non-convex. Biophysics introduces nonlinear thresholds and “cliffs” (fold transitions, aggregation nucleation, chromophore formation), and affinity-improving mutations can destabilize or reduce solubility/specificity [Rabia et al., 2018]. This makes *linear* scalarization of multiple objectives prone to *scalarization bias*: weighted sums recover only supported solutions and miss Pareto-optimal designs in non-convex regions by construction [Van Moffaert et al., 2013, Lin et al., 2024], reducing coverage precisely where biologically viable compromises may lie. While evolutionary methods such as NSGA-II and MosPro can *approximate* the frontier without weighted sums [Deb et al., 2002, Hong and Kortemme, 2024, Luo et al., 2025], their iterative sampling can be inefficient in large sequence spaces and offer no trade-off controllability.

Pretrained protein language models (PLMs) provide strong priors over plausible sequences [Ferruz et al., 2022, Nijkamp et al., 2023]. When design preferences shift, inference-time controllability is preferable to retraining. Inference-time alignment via autoregressive reward models (ARMs) steers a frozen base model efficiently (including multi-objective trade-offs as in GenARM and PARM [Xu et al., 2025, Lin et al., 2025a]) but common instantiations still aggregate objectives linearly and assume deterministic winner–loser supervision from noisy evaluators, leading to *uncertainty-blind* pairwise learning.

We present Smooth Tchebycheff Preference-Aware Reward Model (ST-PARM), an inference-time alignment framework for multi-objective protein design. It retains a frozen PLM prior while learning a small autoregressive reward model, cross-objective trade-off preference-conditioned, to steer generation at inference time. In our protein studies, a ∼10^6^-parameter reward model steers a frozen, ∼10^9^-parameter base model.

Within this novel framework for protein design, ST-PARM makes two major contributions. First, it addresses scalarization bias by replacing linear aggregation with a smooth Tchebycheff scalarization that is Pareto-complete in principle. Second, it addresses uncertainty-blind preference learning by introducing a reward-calibrated pairwise loss that down-weights ambiguous comparisons. It also introduces pair-construction strategies based on latent-space clustering to form informative within-and across-cluster comparisons.

We evaluate ST-PARM on GFP fluorescence–stability (with measured fluorescence [Sarkisyan et al., 2016]) and IL-6 CDR3+suffix stability–solubility, improving Pareto coverage and tradeoff preference tracking over PARM and MosPro in the unfiltered setting. On GFP, a conservative structural screen (local confidence and global fold preservation) retains a broad frontier with post-filter coverage/controllability close to unfiltered values. We also report cross-evaluator checks, a three-objective extension, and a natural-language alignment generality check (Supplement).

Figure 1 provides a graphical overview of ST-PARM, summarizing the targeted gaps, the corresponding fixes (reward-calibrated pairwise preference and smooth Tchebycheff scalarization), and the evaluated instantiations and outcomes.

**Fig 1.**
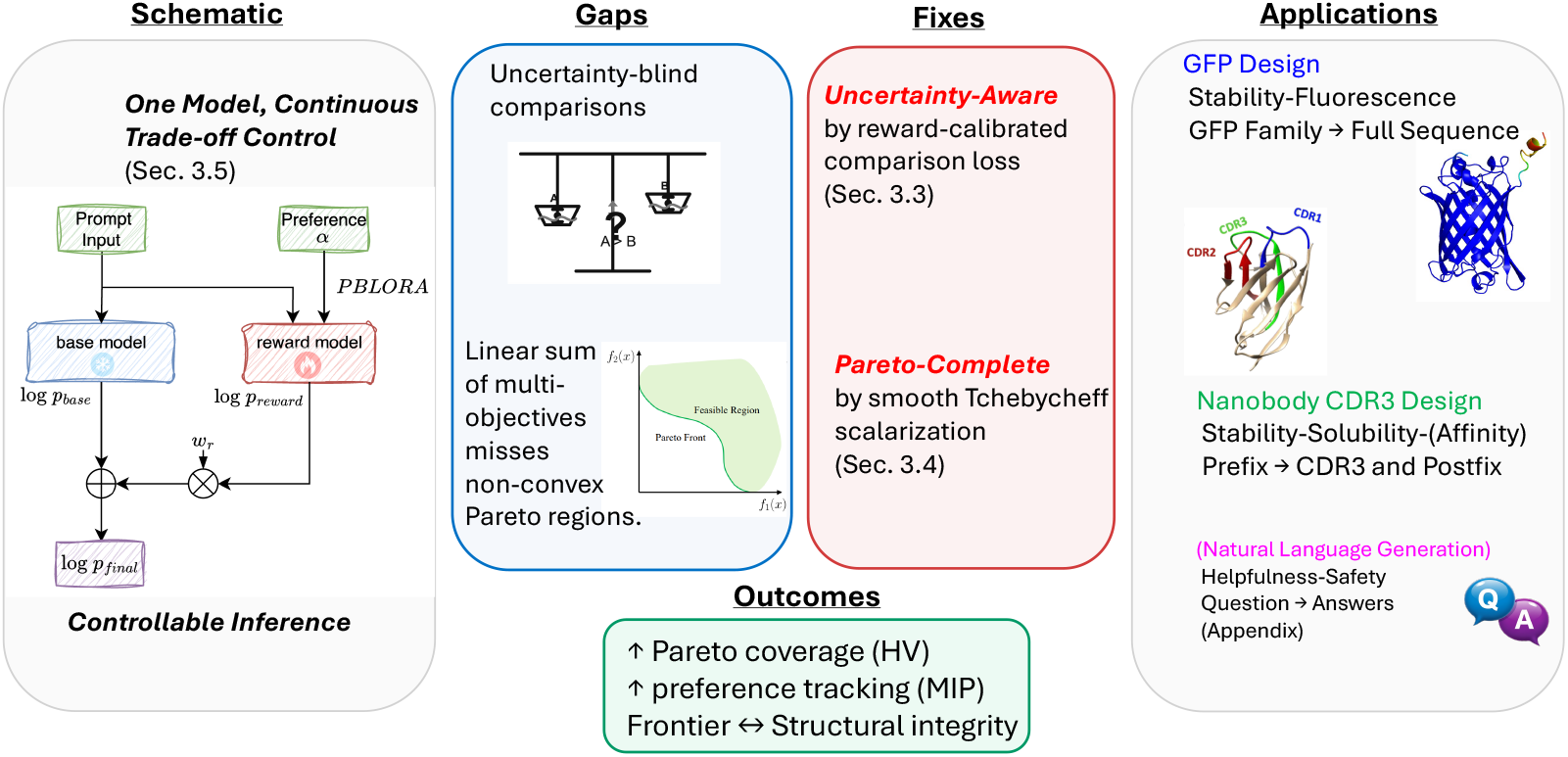
ST-PARM overview. A frozen base model is steered at inference time by a preference-conditioned ARM (input trade-off *α*). Reward calibration addresses uncertainty-blind comparisons; smooth Tchebycheff mitigates linear-scalarization bias, enabling one-model controllability and improved Pareto coverage (HV) and tradeoff preference tracking/controllability (MIP).

## 2. Methods

### 2.1. Problem Setup and Notation

Let 𝒳 denote the space of prompts (e.g., protein scaffolds, sequence prefixes, or natural language queries) and 𝒴 the space of candidate designs (e.g., amino acid sequences), also known as responses (e.g. natural text answers). Both prompt *x* ∈ 𝒳 and response *y* are a sequence of tokens (e.g., amino acids for protein sequences or words for natural texts), for which *y*_*t*_ denotes the *t*-th token of *y* and *y*_<*t*_ the prefix of *y*_*t*_.

We assume *k* objectives when optimizing response *y*. For objective *i* ∈ {1, …, *k*}, an evaluator *f*_*i*_(·), computational or experimental, provides a continuous, scalar label that defines the preference between paired responses. A trade-off preference vector in the (*k* − 1)-simplex, *α* ∈ Δ^*k*−1^, specifies the desired multi-objective trade-off.

### 2.2. Overview

To enable inference-time controllable multi-objective generation, we learn a *single α*-conditioned ARM *π*_*r*_(· | *x, y*_<*t*_, *α*) to steer a frozen base model. For prompt *x* and response *y*, the ARM defines a token-factorized reward

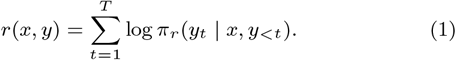

Details and background on ARMs are provided in Supp. A.1. We target two gaps of prior inference-time alignment: *uncertainty-blind* pairwise learning and *linear-scalarization bias*.

### 2.3. Contribution 1: Reward-Calibrated Preference Loss for Uncertainty-Aware Learning

Given continuous, noisy labels *f* for paired (“winning” and “losing”) sequences (*y*^W^, *y*^L^), we replace the standard, deterministic Bradley–Terry training [Bradley and Terry, 1952], as used in modern preference alignment [Rafailov et al., 2023], with a confidence-weighted loss

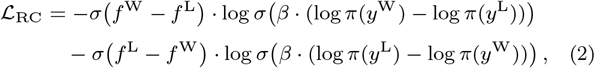

which is equivalent to a cross-entropy between the label-implied preference *p*_*f*_ = *σ*(*f* ^W^ − *f* ^L^) and the policy-implied preference *p*_*π*_ = *σ* (*β* · (log *π*(*y* ^W^) − log *π*(*y* ^L^))): ℒ_RC_ = CE(*p*_*f*_, *p*_*π*_). This yields symmetric, confidence-weighted, noise-robust gradients. Details are in Supp. A.2.

We also cluster sequences in the latent space and introduce within-and cross-cluster pairing strategies besides commonly used random pairing. See details in Supp. B.

### 2.4. Contribution 2: Smooth Tchebycheff Scalarization for Pareto-Complete Learning

Let 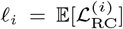 be per-objective calibrated losses. We replace linear scalarization with smooth Tchebycheff scalarization Lin et al. [2024]

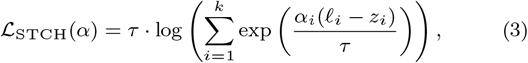

which approaches hard Tchebycheff as *τ* → 0 [Lin et al., 2024] and is Pareto-complete [Lin et al., 2025b]. Details and gradient behavior are in Supp. A.3. We train by sampling *α* ∼ Δ^*k*−1^ and minimizing 𝔼_*α*_[ℒ_STCH_(*α*)].

### 2.5. Trade-off Conditioning for Controllable Inference

We condition a *single* ARM on the user-specified trade-off vector *α* using a parameter-efficient adapter scheme, Preference-aware Bilinear Low-Rank Adaptation (PBLoRA) [Lin et al., 2025a]. Concretely, low-rank adapters are made a smooth function of *α*, with a shared, preference-agnostic component capturing label features common to all objectives and a small preference-aware component steering generation along the requested trade-off. This design keeps memory and compute efficient (we train one ∼10^6^-parameter ARM once) while enabling *continuous* interpolation over *α* at test time. The preference-conditioned ARM then guides decoding with

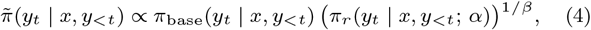

so varying *α* navigates trade-offs without retraining. This stands in contrast to approaches that train a separate model per objective (or per trade-off) as in multi-ARM GenARM-style setups [Xu et al., 2025]. Details are provided in Supp. A.4 and A.5.

### 2.6. Protein Design Benchmarks

ST-PARM is applied to protein design by specifying (i) the prompt *x* (family/scaffold description or sequence prefix), (ii) the design region (full-length vs. CDR3+suffix), and (iii) objective evaluators (computational or experimental) that provide continuous scores for pairwise preference training and for downstream assessment. Table 1 summarizes the two protein benchmarks used in this work.

**Table 1.** Protein-design benchmarks. Full evaluator/backbone details are provided in Supplementary Table S1.

| Protein | Design region | Objectives |
| --- | --- | --- |
| GFP | Full-length | Fluorescence–Stability |
| IL-6 nanobody | CDR3 + suffix | Stability–Solubility( + Affinity in Supp.) |

#### GFP fluorescence–stability

We use an experimentally grounded fluorescence landscape of 51,715 GFP variants [Sarkisyan et al., 2016] and pair it with a sequence-based stability evaluator to form a two-objective benchmark. To mirror low-fitness optimization settings as in MosPro, preference training pairs are constructed (Supp. B) from a bottom-40% filtered subset of 9,769 sequences, leading to 154,944 training pairs and 1,360 validation pairs (Supp. F).

#### IL-6 nanobody CDR3 design

We use a curated IL-6 binding nanobody dataset [Tsuruta et al., 2023] and perform conditional generation of the CDR3 region and suffix from a prefix prompt. Following [Barton et al., 2024], we removed low-germline-identity sequences (< 75%), de-duplicated, and retained IL-6 binders only, yielding 655/81/82 train/val/test sequences. More details on data curation and pair construction are reported in Supp. G. Two-objective results (stability–solubility) are reported in the main text; a three-objective extension is provided in Supp. G.3.

### 2.7. Training and Hyperparameters

Hyperparameters are selected via small-scale grid searches on validation performance, with search ranges and selected values reported in Supp. C. Training uses AdamW with mixed precision (`bf16`); batching, generation counts, and compute details are summarized in Supp. D. Unless otherwise stated, we use the same optimizer schedule and batching protocol across protein tasks and vary only task-specific components (pair construction, prompt format, and evaluators), as detailed in Supp. F and Supplement G. We set *τ* =0.5 and sample *α* ∼ Beta(0.5, 0.5) during training; sensitivity analyses are in Supp. H.

### 2.8. Sequence Generation and Objective Evaluation

At inference time, we generate designs by guided decoding with a frozen base model and a preference-conditioned ARM (Eq. 4). For each trade-off vector *α*, we generate a batch of sequences and evaluate designs with objective-specific evaluators (Table 1): for GFP, fluorescence is assessed with GGS [Kirjner et al., 2024] (trained on Sarkisyan et al., 2016) and stability with TemBERTure [Rodella et al., 2024]; for IL-6 nanobodies, stability and solubility are assessed with DeepSTABp [Jung et al., 2023] and CamSol [Sormanni et al., 2015], respectively, with alternative evaluators (TEMPRO [Alvarez and Dean, 2024]/TANGO [Fernandez-Escamilla et al., 2004]) used for robustness. More details can be found in Supp. F and Supp. G.

On a single NVIDIA A100 (40GB), we observe average generation times of 1.9s/sequence for IL-6 CDR3+suffix and 2.7s/sequence for the full-length GFP. More details are in Supp. D.

### 2.9. Assessment

#### Pareto coverage and preference tracking

We evaluate multi-objective generation using Hypervolume (HV) for Pareto coverage and Mean Inner Product (MIP) for alignment between min-max normalized objective scores of designs and the corresponding objective trade-off vectors. HV is a common assessment metric for Pareto coverage in multi-objective language tasks [Lin et al., 2025a] and protein design [Hong and Kortemme, 2024]. Formal definitions, normalization, and reference-point choices are provided in Supp. E.

#### Protein-specific validation and diagnostics

We treat GFP as the primary testbed for our *generation–validation pipeline*. After sequence generation, we apply a structural validation filter based on AlphaFold2 (v2.3.2; monomer) predictions: average pLDDT ⩾80 and TM-score (to avGFP) ⩾0.5, to obtain the *post-filter* (actionable) cohort. On the post-filter cohort, and for representative designs sampled along the Pareto frontier, we report: (a) sequence-level evaluations—*novelty* (percent identity to the nearest neighbor in the training set and to the wild-type) and *diversity* (portions of 95%-identity clusters); and (b) structure-level evaluations—TM-scores, *C α* RMSD over the TM-aligned region, and chromophore-containing helix RMSD. We also report per-design mean pLDDT and TM-scores prior to filtering to quantify the filter’s effect. Additional details and visualizations are provided in Supp. 3.1.2.

For IL-6 nanobody design, we do not apply the structural filter considering the limited accuracy of nanobody structure prediction. To maintain fair and reproducible scope, IL-6 results are therefore reported on the *pre-filter* cohort only.

## 3. Results and Discussion

We evaluate ST-PARM on two protein-design benchmarks (GFP fluorescence–stability and IL-6 nanobody CDR3+suffix stability– solubility) and report improvements in Pareto coverage (HV) and trade-off controllability (MIP) relative to baselines. For GFP design, we additionally assess sequence profiles and structural integrity on Pareto frontiers. As a generality check beyond proteins, we also report a two-objective natural-language alignment study in Supp. H.

### 3.1. GFP Design on Fluorescence–Stability Benchmark

We assess ST-PARM on the green fluorescent protein (GFP) fluorescence–stability benchmark to test multi-objective optimization with an experimentally grounded objective: fluorescence is measured on a large mutational landscape rather than predicted. This setting provides a stringent sequence–function evaluation and enables direct comparison with MosPro [Luo et al., 2025], which performs mutation-based multi-objective GFP optimization.

We ask whether ST-PARM can generate trade-off preference-controlled designs with improved Pareto coverage and stronger fluorescence–stability trade-offs than prior baselines, including PARM (methodologically closest baseline) and MosPro (recent multi-objective protein-design baseline). We then interpret sequence profiles and structural consequences along the frontier generated by ST-PARM.

*Note*. Cross-method results in Sec. 3.1.1 are unfiltered/pre-filtered (no structural screening). Structural screening is reported for ST-PARM only (Sec. 3.1.2) because baseline structures were unavailable at analysis time.

#### 3.1.1. Frontier Coverage and Trade-off Controllability

##### Frontier visualization over trade-off sweep

Figure 2 summarizes GFP fluorescence–stability trade-offs under trade-off preference conditioning (points colored by *α*_fluor_) and compares ST-PARM against the two baselines, PARM (ablated ST-PARM without the two major contributions) and MosPro.

**Fig 2.**
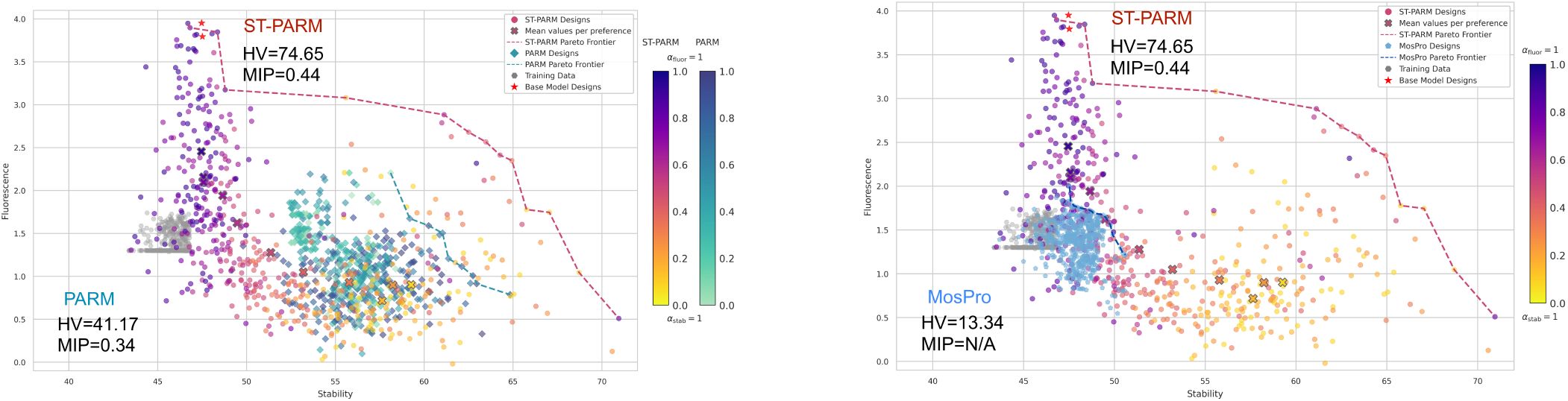
GFP fluorescence–stability trade-offs and Pareto frontier comparison (pre-filter). Left: (A) ST-PARM vs PARM (closest preference-conditioned inference-time alignment baseline). Right: (B) ST-PARM vs MosPro (Pareto protein-design baseline without preference control). Each point is a generated sequence colored by the preference toward fluorescence *α*_fluor_; dashed curves denote aggregated Pareto frontiers computed over all preferences; gray points are training sequences. The colorbar indicates the trade-off direction from fluorescence (*α*_fluor_=1) to stability (*α*_stab_=1).

Across *α*, ST-PARM produces a broader design distribution and an aggregated frontier that extends toward higher fluorescence and stability relative to PARM and MosPro (Fig. 2). Compared to ST-PARM, the PARM designs contract into a narrower band with under-coverage near balanced trade-offs, and the MosPro designs focus in regions close to the training examples. MosPro performs iterative local search by mutation and selection (top 500 among ∼18,000 samples), whereas ST-PARM generates 550 designs (50 per *α*) in a single forward pass conditioned on trade-off preference *α*. Despite being offline, ST-PARM attains a frontier that dominates MosPro across much of the objective space.

Quantitative analyses for Pareto coverage (HV) and tradeoff controllability (MIP) align with the visualization. ST-PARM achieves HV= 74.65 versus PARM HV= 41.17 and MosPro HV= 13.34. For completeness, the mean*±*sd preference-sweep curves are reported in Fig. S5. ST-PARM’s MIP score was 0.44, outperforming PARM’s MIP of 0.35, whereas MosPro does not condition on tradeoff *α* and does not offer such controllability.

Notably, following the benchmark protocol, preference training pairs are constructed from the bottom 40% of sequences in both fluorescence and predicted stability, simulating a low-fitness regime where only weak variants are available. ST-PARM therefore improves trade-offs without direct exposure to high-fluorescence or high-stability examples during training.

##### Ablation study

To isolate the role of the base-model prior, we ablate the base-model prior by generating with the trained reward model alone (i.e., without the GFP-adapted ProLLaMA prior). The resulting Pareto frontier becomes markedly narrower and shifts toward less favorable fluorescence–stability trade-offs (Fig. S8), indicating reduced coverage and degraded optimization. This supports the role of the frozen base model as a biological prior and motivates inference-time alignment as *reweighting* a plausible sequence distribution rather than optimizing a reward model in isolation. Collectively, these results indicate that combining reward-calibrated preference learning with Pareto-complete scalarization improves both frontier coverage and controllable trade-off exploration.

#### 3.1.2. Sequence and Structure Analysis

##### Structural filtering and filtered Pareto

To favor locally confident, fold-preserving designs while preventing proxy exploitation, we apply a conservative structural screen (mean pLDDT ⩾80; TM-score ⩾0.5 vs. WT) and examine the retained set for ST-PARM across *α*. Post-filter set quality remains strong in Pareto coverage (HV = 68.71; pre-filter: = 74.65) and in tradeoff controllability (MIP = 0.45; pre-filter: = 0.44), preserving most coverage/tracking established in the unfiltered analysis (see Supp. Fig. S6 for pre–post 2D overlays).

##### Sequence and structure quality (pre/post)

Table 2 reports *pre-filter* sequence measures for ST-PARM, PARM, and MosPro. It also reports *post-filter* sequence measures for retained ST-PARM designs. ST-PARM sequences maintained superior novelty and diversity compared to MosPro, after quality-conditioned by the structural screen, reflecting useful variety in the final design set. Specifically, the post-filter ST-PARM designs are not near training sequences (96.7% with nearest-neighbor sequence identity [NN seq. id.] below 99%), novel (38.7% with NN seq. id. below 95%, possessing 10+ mutations for GFP of length*>*200), and diverse (95%-cluster count being 54% designs).

**Table 2.** Summary of sequence quality for designs from ST-PARM, PARM, and MosPro. ST-PARM retained design quality under structural screening.

| Model | Novelty (NN seq. id.) |  |  | Diversity<br>95%-cluster |
| --- | --- | --- | --- | --- |
| | median | $\geq 99\%$ | $\leq 95\%$ | |
| MosPro | 99.6% | 98.4% | 0.8% | 4.0% |
| PARM | 51.6% | 0.0% | 100% | 74.7% |
| ST-PARM | 91.6% | 1.9% | 59.6% | 69.6% |
| ST-PARM (filtered) | 96.2% | 3.3% | 38.2% | 54.0% |

Additional structural measures further support the quality of the filtered ST-PARM designs. Their median pLDDT and TM scores are 95.9 and 0.99, respectively. 81.9% are with TM scores above 0.8, reflecting highly preserved functional folds. Baseline structural predictions were unavailable at the time of analysis, so structural metrics are not compared head-to-head.

##### Interpretation along the retained frontier

For the retained ST-PARM designs, we select representative designs along the fluorescence–stability Pareto frontier and report sequence identity and three structural similarity diagnostics relative to the wild-type avGFP reference: (i) TM score (major diagnostic); (ii) backbone RMSD of TM-aligned region, capturing global *β*-barrel deviations, and (iii) helix RMSD, focusing on chromophore-containing helix (residue 58–71 of avGFP). These quantitative assessment metrics are experimentally validated to assess and prioritize GFP computational designs [Hayes et al., 2025, Zhu et al., 2025]. As shown in Fig. 3, fluorescence-leaning designs generally exhibit lower helix RMSD and higher sequence identity (as seen in a “phase” shift) with high fold preservation (as seen in high TM scores; not monotone due to TM filter’s gating effect and finite sampling), whereas stability-leaning settings introduce larger sequence changes and correspondingly higher helix RMSDs. Changing the TM-score filter from ⩾0.5 to ⩾0.7 or ⩾0.8 leads to similar trends (Supp. Fig. S7).

**Fig 3.**
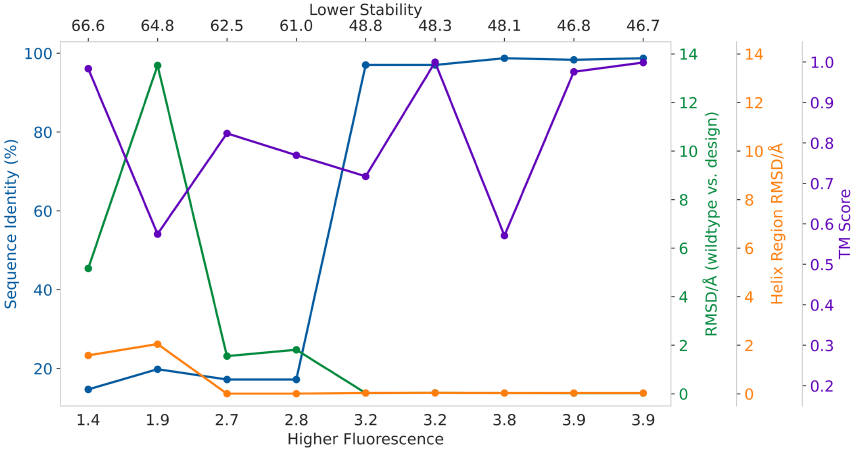
Sequence and structure variation along the GFP fluorescence–stability Pareto frontier (filtered ST-PARM). Reported are sequence identity, TM scores, backbone RMSD (TM-aligned region), and chromophore-containing helix RMSD of representative frontier designs relative to wild-type avGFP, as the preference to fluorescence increases toward the right.

##### Sequence profiles (unfiltered)

We compare sequence logos for (i) natural GFP variants, (ii) the GFP-adapted base model (GFP-finetuned ProLLaMA prior to ARM steering), and (iii) **unfiltered** ST-PARM designs aggregated over the fluorescence–stability frontier (Supplement F.4). The base model largely recapitulates dominant preferences observed in natural GFPs, whereas ST-PARM steering increases per-position diversity while typically preserving the dominant residue at conserved sites.

### 3.2. IL-6 Nanobody CDR3 Design

We next demonstrate generality of ST-PARM on a therapeutically motivated nanobody design task by generating IL-6 nanobody CDR3+suffix sequences conditioned on a CDR3-prefix prompt that is derived from the held-out test set.

#### 3.2.1. Frontier Coverage and Trade-off Controllability

##### Frontier visualization over trade-off sweep

We consider two developability-related objectives—thermodynamic stability (melting-temperature prediction) and solubility (sequence-derived solubility score). For each inference-time trade-off vector *α* = (*α*_sta_, *α*_sol_) ∈ {(0.0, 1.0), (0.1, 0.9), …, (1.0, 0.0)}, we generate sequences conditioned on test-set CDR3 prefixes and evaluate designs using DeepSTABp (stability) and CamSol (solubility). Figure 4 shows a smooth and continuous trade-off frontier as *α* shifts the preference from solubility to stability, indicating controllable multi-objective generation.

**Fig 4.**
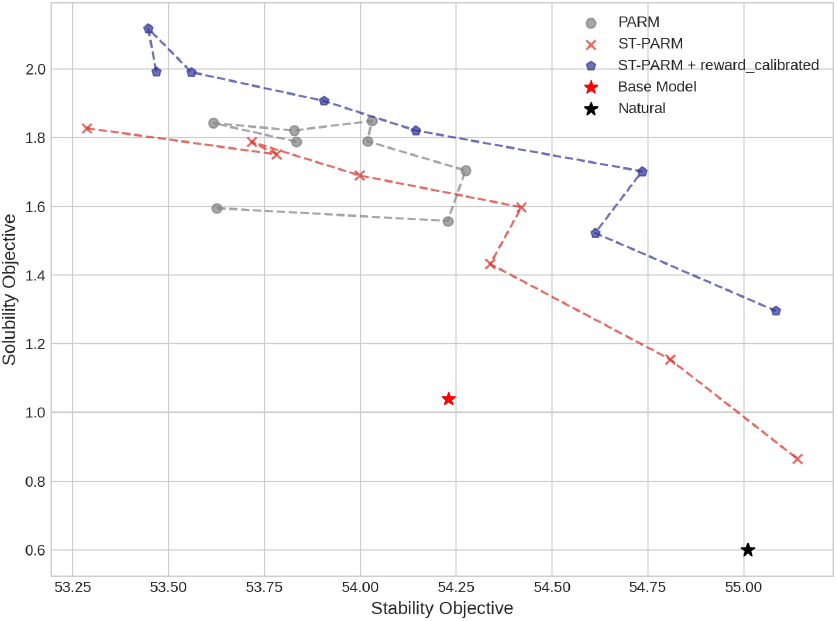
IL-6 nanobody design: stability–solubility trade-off under inference-time preferences. Each point is the mean predicted stability and solubility of sequences generated under a trade-off vector *α* = (*α*_sta_, *α*_sol_). As *α* shifts the preference from solubility to stability, ST-PARM traces a smooth trade-off curve, indicating continuous controllability.

##### Quantitative evaluation and ablation

Table 3 reports HV (Pareto coverage) and MIP (preference tracking) for PARM and ST-PARM, with and without reward-calibrated preference learning. ST-PARM with Pareto-complete scalarization alone improves both HV and MIP, and reward calibration further increases coverage and preference adherence.

**Table 3.**
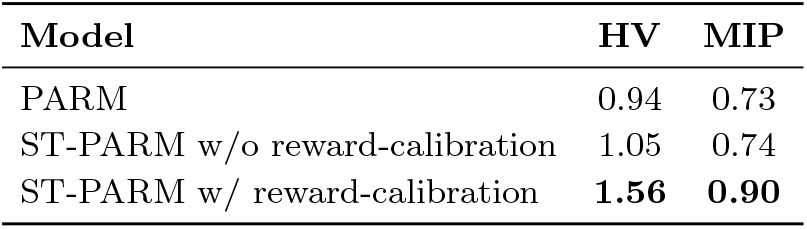
IL-6 nanobody two-objective quantitative summary and ablations. HV measures Pareto coverage and MIP measures trade-off preference tracking across the inference-time preference sweep.

| Model | HV | MIP |
| --- | --- | --- |
| PARM | 0.94 | 0.73 |
| ST-PARM w/o reward-calibration | 1.05 | 0.74 |
| ST-PARM w/ reward-calibration | **1.56** | **0.90** |

##### Robustness and higher-dimensional extension (Supplement)

We further extend IL-6 design to three objectives (stability– solubility–affinity); results are in Supp. G.3. We additionally re-score generated designs using alternative predictors (TEMPRO for stability and TANGO for solubility) and observe positive rank agreement (Supp. G.4).

## 4. Conclusion

We presented ST-PARM, a framework for trade-off preference-conditioned, multi-objective sequence generation that addresses key limitations of conventional pairwise preference learning and linear scalarization. ST-PARM combines a reward-calibrated preference loss that down-weights uncertain comparisons with a smooth Tchebycheff scalarization that improves coverage of non-convex trade-off regions. Together with parameter-efficient trade-off preference conditioning, ST-PARM enables inference-time controllability over trade-offs without retraining the reward model. The framework is lightweight in computation, training a small reward model (∼ 10^6^ parameters in our protein studies) once to steer a much larger frozen generator (∼ 10^9^ parameters).

Across GFP fluorescence–stability and IL-6 nanobody CDR3+suffix design, ST-PARM expands Pareto coverage and improves trade-off controllability against PARM and MosPro, as summarized by frontier visualizations and HV/MIP quantification. For GFP it preserves most of that coverage after a conservative structural screen, yielding an actionable retained set for downstream experiments. For IL-6, evaluator-robustness checks and three-objective scalability are provided in the Supplement.

A key limitation is that, while GFP fluorescence supervision is experimentally grounded, evaluations of generated sequences for stability, solubility, and affinity rely on computational predictors. Future work should incorporate prospective wet-lab validation. In parallel, integrating structure-aware objectives or priors during generation may further increase retained coverage under stricter TM filters.

By unifying uncertainty-aware preference learning, Pareto-complete scalarization, and preference conditioning, ST-PARM provides a practical foundation for controllable sequence generation under competing objectives and noisy evaluators.

## 6. Author contributions statement

R.Y. and Y.S. designed the study and conceived the experiments. R.Y. conducted the experiments. R.Y. and Y.S. analyzed the results. R.Y. and Y.S. wrote and reviewed the manuscript.

## 7. Acknowledgments

This work is in part supported by the National Institute of General Medical Sciences (R35GM124952 to Y.S.) and the National Science Foundation (1943008 to Y.S.). Computing resources were provided in part by High Performance Research Computing at Texas A&M University.

LLM use disclosure: Portions of the manuscript text were edited for language and readability using GPT-5.2 as an editorial aid. The authors wrote the original draft and reviewed, verified, and revised all LLM-suggested edits.

