## Supplementary material for "ST-PARM: Pareto-Complete Inference-Time Alignment for Multi-Objective Protein Design": Supp.

### A. Supplementary Methods: Core Framework

#### A.1. Autoregressive Reward Modeling (full background)

We solve the multi-objective protein design problem (and, in general, sequence generation) by inference-time alignment. Specifically, during inference time, we steer a frozen base autoregressive generator $\pi_{\text{base}}$ with a lightweight autoregressive reward model (ARM) $\pi_r$. ARMs provide a theoretically principled approach for inference-time alignment of sequence models.

**ARM parameterization and sequence-level reward.** We model reward signals using an autoregressive reward model (ARM) that provides left-to-right token scores $\pi_r(y_t \mid x, y_{<t})$. The sequence-level reward for a response $y$ conditioned on a prompt $x$ is the token-log-likelihood under the ARM:

$$r(x,y) \;=\; \sum_{t=1}^{T} \log \pi_r(y_t \mid x, y_{<t}). \tag{S1}$$

Equivalently, we can write a token-factorized decomposition

$$\begin{aligned} r(x,y) &= \sum_{t=1}^{T} r_t(x, y_t \mid y_{<t})\,, \text{where} \\ r_t(x, y_t \mid y_{<t}) &:= \log \pi_r(y_t \mid x, y_{<t})\,. \end{aligned} \tag{S2}$$

so that each token contributes a reward increment rather than a single scalar at the end of the sequence. This yields *dense credit assignment* and facilitates stable optimization and efficient supervision at the token level.

**Closed-form alignment under KL constraints.** For KL-regularized alignment with inverse temperature $\beta$, the optimal preference-adjusted decoding distribution satisfies

$$\log \pi_{\text{dec}}(y \mid x) \;=\; \log \pi_{\text{base}}(y \mid x) \;+\; \tfrac{1}{\beta}\, r(x,y) \;-\; \log Z(x), \tag{S3}$$

which, using (S2), decomposes autoregressively as

$$\pi_{\text{dec}}(y_t \mid x, y_{<t}) \;\propto\; \pi_{\text{base}}(y_t \mid x, y_{<t}) \;\exp\!\Big(\tfrac{1}{\beta}\, r_t(x, y_t \mid y_{<t})\Big). \tag{S4}$$

Thus, ARMs yield a *closed-form* next-token update rule: alignment reduces to an additive shift of base-model logits by reward-model logits, eliminating the need for on-policy RL and the instabilities associated with policy optimization.

**Benefits of autoregressive factorization.** Equation (S4) provides several advantages:

- **Local credit assignment.** Each token receives an explicit, differentiable reward increment $r_t$, improving learnability relative to trajectory-only scalar rewards.
- **Compatibility with base architectures.** Because $\pi_{\text{base}}$ and $\pi_r$ share the same autoregressive structure, alignment can be implemented by modifying logits at test time without retraining the base model.
- **Stable optimization.** The KL-regularized decoding rule arises analytically, avoiding the variance and divergence issues typical of reinforcement learning methods.

#### A.2. Reward-Calibrated Preference Loss: full derivation and properties

Pairwise preference learning.

Given a dataset of prompt–response tuples $(x, y^{\mathrm{W}}, y^{\mathrm{L}})$, where the "winning" response $y^{\mathrm{W}}$ is superior to the "losing" $y^{\mathrm{L}}$, common ARMs are trained via a Bradley–Terry style logistic loss:

$$\mathcal{L}_{\text{pair}} = -\log \sigma\left(\beta \cdot (r(x, y^{\mathrm{W}}) - r(x, y^{\mathrm{L}}))\right), \tag{S5}$$

where $\sigma(\cdot)$ is the logistic sigmoid and $\beta > 0$ is a margin-scaling hyperparameter.

Standard loss for pairwise preference training (Eq. S5) treats all pairwise comparisons equally certain. In scientific design, objective evaluations or labels (including computational predictions and experimental measurements), such as molecular properties, are often continuous yet noisy. Many pairwise comparisons are actually near ties within evaluator uncertainty. To address this challenge, we propose a *reward-calibrated pairwise preference loss*, which directly incorporates the magnitude and direction of the label difference to calibrate the reward-based loss. Given single-objective labels $f^{\mathrm{W}}$ and $f^{\mathrm{L}}$ for sequences $y^{\mathrm{W}}$ and $y^{\mathrm{L}}$, and corresponding log-probabilities (policy scores) $\log \pi(y^{\mathrm{W}})$ and $\log \pi(y^{\mathrm{L}})$, the loss for a single pair is defined as:

$$\begin{aligned} \mathcal{L}_{\text{RC}} = &-\sigma\big(f^{\mathrm{W}} - f^{\mathrm{L}}\big) \cdot \log \sigma\big(\beta \cdot (\log \pi(y^{\mathrm{W}}) - \log \pi(y^{\mathrm{L}}))\big) \\ &- \sigma\big(f^{\mathrm{L}} - f^{\mathrm{W}}\big) \cdot \log \sigma\big(\beta \cdot (\log \pi(y^{\mathrm{L}}) - \log \pi(y^{\mathrm{W}}))\big) \end{aligned} \tag{S6}$$

where $\sigma(\cdot)$ is the logistic sigmoid, $\beta > 0$ controls the sharpness of the policy margin. When the policy $\pi$ is represented by a learned autoregressive reward model $\pi_r(y_t \mid x, y_{<t})$, the log-probability terms $\log \pi(y)$ can be decomposed token-wise as in Eq. 1.

This loss structure ensures several desirable properties. (1) Continuity and symmetry. The loss is continuous and symmetric under label permutation. No explicit pairwise preference labeling is needed; preference is inferred from the label gap itself. (2) Confidence-weighted learning. The loss smoothly interpolates between strong preference (large label difference) and abstention (small or ambiguous label difference). (3) Noise robustness. The form penalizes both false positives and false negatives, reducing the impact of stochasticity in objective evaluations.

Probabilistic interpretation.

The reward-calibrated loss in Eq. S6 can be viewed as a cross-entropy between two Bernoulli distributions: a label -implied preference distribution $p_f = \sigma(f^{\mathrm{W}} - f^{\mathrm{L}})$ and a policy-implied preference distribution $p_\pi = \sigma\left(\beta \cdot (\log \pi(y^{\mathrm{W}}) - \log \pi(y^{\mathrm{L}}))\right)$. The loss becomes:

$$\mathcal{L}_{\text{RC}} = \text{CE}(p_f, p_\pi) = -p_f \log p_\pi - (1 - p_f) \log(1 - p_\pi), \tag{S7}$$

where $\text{CE}(p_f, p_\pi)$ denotes the cross-entropy. This interpretation formalizes the idea that the policy $\pi$ should match the probabilistic ordering implied by the label $f$ gap. Rather than enforcing a hard ranking, the model learns to approximate the relative preference structure encoded in label differences, leading to smoother gradients and improved generalization.

Preference pairs.

For protein datasets where sequences are property-labeled but not naturally paired, we construct training pairs with three strategies. The common strategy is random pairing. We additionally propose and test embedding-based pairing strategies—within-cluster (local comparisons) and across-cluster (global comparisons) in the latent space—and select the best-performing strategy per dataset. We report pairing details and ablations in Supp. B and F.

##### A.3. Smooth Tchebycheff scalarization: Pareto completeness and gradients

**Multi-objective extension (GenARM-style).** For $k$ objectives with $k$ ARMs $\{\pi_r^{(i)}\}$ $(i = 1, \ldots, k)$ and trade-off preference vector (or tradeoff vector in short) $\alpha \in \Delta^{k-1}$, the composite reward $r_\alpha = \sum_{i=1}^{k} \alpha_i r^{(i)}$ yields the decoding rule

$$\pi_\alpha(y_t \mid x, y_{<t}) \;\propto\; \pi_{\text{base}}(y_t \mid x, y_{<t}) \prod_{i=1}^{k} \exp\Big(\tfrac{\alpha_i}{\beta}\, r_t^{(i)}(x, y_t \mid y_{<t})\Big). \tag{S8}$$

This multiplicative form shows that each objective contributes an additive logit shift weighted by the user preference $\alpha$. Theoretically, this guarantees:

- smooth interpolation between objectives,
- coherent movement along Pareto frontiers, and
- explicit control of trade-offs at the token level.

**Preference-conditioned ARMs (PARM-style).** Instead of maintaining $k$ separate ARMs, PARM parameterizes a *single* preference-conditioned ARM $\pi_r(\cdot \mid x, y_{<t}, \alpha)$ with reward

$$r_\alpha(x, y) \;=\; \sum_{t=1}^{T} \log \pi_r(y_t \mid x, y_{<t}, \alpha). \tag{S9}$$

The corresponding decoding distribution becomes

$$\pi_\alpha(y_t \mid x, y_{<t}) \;\propto\; \pi_{\text{base}}(y_t \mid x, y_{<t}) \big[\pi_r(y_t \mid x, y_{<t}, \alpha)\big]^{1/\beta}, \tag{S10}$$

which unifies multi-objective alignment into a single network conditioned on $\alpha$. This provides:

- changes in $\alpha$ induce continuous changes in $\pi_\alpha$;
- objectives inform each other through a shared representation;
- no re-training is needed to evaluate new preferences.

**ST-PARM.** In multi-objective settings, for objective $i$ we define a reward-calibrated per-objective loss for a batch of training tuples by:

$$\ell_i = \mathbb{E}_{(x, y^{\mathrm{W}}, y^{\mathrm{L}})}\left[\mathcal{L}_{\mathrm{RC}}^{(i)}(x, y^{\mathrm{W}}, y^{\mathrm{L}})\right], \tag{S11}$$

Previous frameworks such as PARM aggregate (not reward-calibrated) objectives via linear scalarization: $\mathcal{L}_{\text{linear}} = \sum_{i=1}^{k} \alpha_i \ell_i$, where $\alpha \in \Delta^{k-1}$ is a trade-off preference vector.

To mitigate scalarization bias and improve coverage of non-convex trade-off regions, ST-PARM uses Smooth Tchebycheff (STCH) scalarization.

$$\mathcal{L}_{\text{STCH}}(\alpha) = \tau \cdot \log\left(\sum_{i=1}^{k} \exp\left(\frac{\alpha_i(\ell_i - z_i)}{\tau}\right)\right), \tag{S12}$$

where $z_i$ is a reference point (typically chosen as the ideal or minimum achievable value of objective $i$; in our use case, we use 0 for all $z_i$ as the sub-losses $l_i$ are always non-negative.), and $\tau > 0$ is a temperature parameter controlling the smoothness of the approximation.

Pareto Completeness.

As $\tau \to 0$, Eq. S12 converges pointwise to the *hard* (or classical) Tchebycheff scalarization: $\lim_{\tau \to 0} \mathcal{L}_{\text{STCH}}(\alpha) = \max_i \alpha_i(\ell_i - z_i)$, which is known to be *Pareto-complete* [Lin et al., 2025b]. The smooth variant in Eq. S12 preserves differentiability, and sampling $\alpha$ during training enables coverage across trade-off regions of the Pareto frontier.

Gradient behavior.

The gradient of the STCH objective softly prioritizes the most underperforming objective relative to the current preference. When $\tau$ is small, the scalarized loss behaves similarly to the maximum of weighted deviations $\alpha_i(\ell_i - z_i)$, concentrating updates on the most violated constraint. As $\tau$ increases, the scalarization becomes more uniform, distributing learning signal across all objectives. This gives rise to a flexible trade-off between focused correction and global convergence.

##### A.4. Preference conditioning via PBLoRA

To enable inference-time controllability over arbitrary trade-off preference vectors $\alpha \in \Delta^{k-1}$, we adopt a parameter-efficient conditioning mechanism based on Preference-aware Bilinear Low-Rank Adaptation (PBLoRA). We follow PARM to construct a single unified autoregressive reward model $\pi_r(y \mid x; \alpha)$ whose weights are modulated by $\alpha$.

Let $\theta_0$ denote the (frozen) parameters of the pretrained base model. The trade-off preference-conditioned parameters $\theta(\alpha)$ of the reward model are computed as: $\theta(\alpha) = \theta_0 + s \cdot BW(\alpha)A$, where $B, A$ are low-rank projection matrices, $W(\alpha)$ is an $\mathbb{R}^{r \times r}$ matrix computed from $\alpha$ via a small neural network, and $s$ is a scaling factor. The bilinear form $BW(\alpha)A$ enables expressive modulation with $\mathcal{O}(r^2)$ additional parameters.

We decompose the update into preference-agnostic and preference-aware terms: $BW(\alpha)A = B_1 W_1 A_1 + B_2 W_2(\alpha) A_2$. The first term captures shared label features across objectives, while the second specializes the model to user-specified trade-offs.

Training Objective

The final training objective is the expected smooth Tchebycheff scalarized loss over sampled trade-off preference vectors:

$$\min_{\Theta} \mathbb{E}_\alpha \left[\mathcal{L}_{\text{STCH}}(\alpha)\right], \tag{S13}$$

where $\Theta$ denotes the PBLoRA parameters, and each $\mathcal{L}_{\text{STCH}}(\alpha)$ is computed from the calibrated per-objective losses $\ell_i$ using Eq. S12. During each training step, we sample $\alpha$, compute $\theta(\alpha)$ via PBLoRA, evaluate $\ell_i$ using Eq. 2, and update $\Theta$ by gradient descent.

##### A.5. Inference-time decoding

Once trained, the ST-PARM model enables controllable sequence generation by modulating the frozen base model $\pi_{\text{base}}$ using the trained, preference-conditioned ARM $\pi_r(y \mid x; \alpha)$. At inference time, we compute a modified next-token distribution:

$$\tilde{\pi}(y_t \mid x, y_{<t}) \propto \pi_{\text{base}}(y_t \mid x, y_{<t}) \cdot (\pi_r(y_t \mid x, y_{<t}; \alpha))^{1/\beta}, \tag{S14}$$

where $\beta$ is a temperature parameter controlling the strength of alignment guidance. As $\beta \to \infty$, generation reverts to the base model; as $\beta \to 0$, alignment becomes more aggressive.

By varying the input trade-off preference vector $\alpha$, users can explore diverse trade-offs on the learned Pareto frontier without retraining or fine-tuning. This facilitates flexible design exploration in domains such as antibody engineering, where objectives like affinity, solubility, and stability must be simultaneously optimized under evolving application constraints.

#### B. Supplementary Methods: Pairing Strategy

To construct a paired dataset from the original property-labeled, unpaired protein sequences, we first obtain fixed-length representations of all sequences using a LM sequence encoder, followed by a nonlinear dimensionality reduction to project them into a low-dimensional embedding space. The reduced embeddings provide a compact representation of sequence space that preserves both local similarity and global diversity among variants. We then apply clustering to this representation to partition the sequences into groups of similar structural or functional characteristics.

Based on these clustered embeddings, we design three distinct strategies for forming sequence pairs. Each strategy emphasizes a different level of sequence diversity and property contrast, and is applied to examine its effect on model learning. When the number of possible pairs is computationally manageable, all pairs resulted are used for training.

- **Across-cluster pairing.** Sequence pairs are sampled between distinct clusters to emphasize global diversity. This strategy encourages the model to learn broad distinctions in the sequence-property relationship, exposing it to pairs that likely differ substantially in sequence composition and biophysical characteristics.
- **Within-cluster pairing.** Sequence pairs are sampled within the same cluster to focus on fine-grained variations among closely related sequences. To ensure informative preference signals, we exclude pairs whose property values are too similar, thereby avoiding nearly indistinguishable comparisons that provide weak supervision. This filtering retains pairs that reflect nontrivial but still localized differences in biophysical properties, allowing the model to capture subtle sequence-property dependencies within a coherent structural context.
- **Random pairing.** Sequence pairs are sampled uniformly at random across the entire dataset, regardless of cluster membership. This serves as a baseline configuration that mixes both local and global relationships without explicit control over sequence similarity or property contrast.

Each pairing scheme represents a different hypothesis about the most informative comparative structure in the data, ranging from global contrasts to fine-grained local differences, allowing systematic assessment of how pair composition influences preference-based learning in protein design.

#### C. Supplementary Methods: Hyperparameter Settings

For all experiments, hyperparameters were selected through small-scale grid searches based on validation performance. Learning rates were explored in the range $\{5\times10^{-4},\, 1\times10^{-4},\, 5\times10^{-5}\}$, and the optimal value for each task was chosen according to the averaged validation accuracy. The number of training epochs was tuned within $\{1,2,3,4,5,6\}$ and similarly selected using validation performance.

The smoothing coefficient $\tau$ and the concentration parameter of the preference distribution Beta(`param`, `param`) were each varied independently as reported in Supp. H, to assess their influence on trade-off controllability and Pareto coverage.

The inverse-temperature parameter $\beta$ in the preference loss was tuned within $\{5\times10^{-2},\, 1\times10^{-2},\, 5\times10^{-3}\}$. For multi-objective experiments, identical $\beta$ values were applied across objectives, as no prior assumption justified asymmetric scaling between reward heads. Within the tested range, model performance was largely insensitive to moderate changes in $\beta$, suggesting stability of optimization dynamics with respect to this parameter.

#### D. Supplementary Methods: Training and Generation

**Hardware usage.** Model training was conducted with 8 NVIDIA A100 GPUs (40GB memory), while sequence generation and evaluation jobs with 4 A100 GPUs.

**Optimizer and schedule.** We trained all models using the AdamW optimizer with a weight decay of 0.05 and an initial learning rate of $5\times10^{-4}$. The learning rate followed a linear warm-up for the first 20 steps, then decayed according to a cosine schedule over 3 training epochs.

**Precision and memory.** To improve computational efficiency and fit larger batches per GPU, we employed bfloat16 mixed precision (`bf16`) training. These settings significantly reduced memory consumption without observable degradation in model accuracy or convergence stability.

**Batching** During training, an effective global batch size of 32 preference (comparison) pairs was maintained across 8 GPUs. Each device processed a per-GPU batch size of 2, with gradient accumulation over 2 steps to achieve the target global batch. During generation, a batch size of 32 sequences was maintained across 4 GPUs, with a per-GPU batch size of 8. The number of sequences generated for each experiment was as follows. Protein family-prompted GFP design: 550; Test CDR3 prefix-prompted IL-6 design: 792 (two-objective) or 4,752 (three-objective); and NLP question-promoted answering: 1,500.

**Training, generation, and evaluation speed.** Each training job on 8 GPUs takes at most 10 hours for two objectives and around 27 hours for three objectives. During inference, each job takes at most 5 hours on 4 A100 GPUs to finish sample generation and property evaluation with oracle models. Specific properties and oracle models are detailed subsequently for each design or generation experiment.

For both protein design tasks, we report average wall-clock generation time per sequence. Per NVIDIA A100 GPU (40GB), IL-6 nanobody design requires approximately `1.9` seconds to generate a sequence. For the GFP fluorescence—stability design task, the average generation cost is `2.7` GPU seconds to generate a sequence.

#### E. Supplementary Methods: Assessment Metrics

We first evaluate multi-objective generation performance using two complementary metrics that are domain-agnostic: the *Hypervolume* (HV) for Pareto coverage and the *Mean Inner Product* (MIP) or trade-off preference alignment.

**HV for Pareto coverage.** Let $\mathcal{S} = \{\mathbf{r}^{(1)}, \mathbf{r}^{(2)}, \ldots, \mathbf{r}^{(N)}\}$ denote the set of per-objective reward vectors obtained from generated samples under varying trade-off preference vectors $\alpha \in \Delta^{k-1}$, where $\mathbf{r}^{(i)} = (r_1^{(i)}, \ldots, r_k^{(i)}) \in \mathbb{R}^k$. The Hypervolume quantifies the volume of the region in the objective space that is dominated by $\mathcal{S}$ with respect to a reference point $\mathbf{z}_{\text{ref}}$, defined as:

$$\mathrm{HV}(\mathcal{S}) = \Lambda\big(\{\mathbf{p} \in \mathbb{R}^k \mid \exists \mathbf{r} \in \mathcal{S},\ \mathbf{r} \preceq \mathbf{p} \preceq \mathbf{z}_{\text{ref}}\}\big), \tag{S15}$$

where $\Lambda(\cdot)$ denotes the Lebesgue measure and $\preceq$ represents elementwise dominance. A higher HV indicates broader Pareto frontier coverage and better diversity of trade-off solutions.

**Table S1.** Protein-design benchmarks used in this work. The ProLLaMA backbone is frozen during ST-PARM training; a lightweight preference-conditioned ARM is trained via PBLoRA. Objective evaluators are used to construct preference pairs and/or to assess generated designs. Additional evaluators are used for robustness checks in Supp..

| Protein | Prompt $x$ | Design $y$ | Backbone / trained ARM | Objectives | Evaluators |
|---|---|---|---|---|---|
| GFP | GFP superfamily | Full-length sequence | ProLLaMA-7B (frozen; GFP-adapted via LoRA) + ARM (PBLoRA; ~6M) | Fluorescence–Stability | Fluorescence: Exp. labels (training) / GGS (reporting) Stability: TemBERTure |
| IL-6 nanobody | CDR3 prefix | CDR3 + suffix | ProLLaMA-7B (frozen) + ARM (PBLoRA; ~6M) | Stability–Solubility (+ Affinity in Supp.) | DeepSTABp–CamSol (TEMPRO–TANGO–BindPred in Supp.) |

**MIP for trade-off preference alignment.** We compute the Mean Inner Product (MIP) after min-max normalizing each objective's scores to the range $[0, 1]$ across all generated samples to remove scale disparities among objectives. Let $\tilde{\mathbf{r}}^{(i)} = (\tilde{r}_1^{(i)}, \ldots, \tilde{r}_k^{(i)})$ denote the normalized label vector for the $i$-th preference setting $\alpha^{(i)}$. The MIP is then defined as:

$$\text{MIP} = \frac{1}{N}\sum_{i=1}^{N}\langle \alpha^{(i)}, \tilde{\mathbf{r}}^{(i)}\rangle, \tag{S16}$$

which measures the directional alignment between the normalized objective scores and the corresponding trade-off preference vectors. A higher MIP indicates that the model's generated outputs more closely follow the intended preference trade-offs. Together, HV and MIP jointly assess the diversity and controllability of the model's multi-objective behavior.

For protein design, we also assess generated sequence profiles and AlphaFold2-predicted structures.We provide the setup including objectives and their evaluators in Table S1 as well as the detailed protocols and results in the subsequent sections.

#### F. GFP Design: Protocols and Supplementary Results

##### F.1. Training Dataset Generation

**Data Sources and Property Definitions.** This benchmark builds on the large-scale experimental fitness landscape of GFP reported by [Sarkisyan et al., 2016], which provides fluorescence measurements for 51,715 GFP variants derived from random mutagenesis of the wild-type *Aequorea victoria* GFP (avGFP) sequence. This dataset constitutes one of the most comprehensive single-gene mutational scans available, capturing both additive and epistatic effects of mutations on fluorescence. Each sequence in the dataset is associated with an experimentally measured fluorescence brightness value, reflecting the functional fitness of that variant in a cellular context. The MosPro framework [Luo et al., 2025] later utilized the same dataset to study the fluorescence–stability trade-off in GFP by computing $\Delta\Delta G$ stability changes for all variants using FoldX, thereby defining a two-property benchmark dataset.

In this work, we adopt the same source data but redefine stability in a way that is consistent with our generative modeling framework. Since our model performs *de novo* sequence generation unconstrained by a fixed reference structure, the FoldX-based $\Delta\Delta G$ values used in MosPro cannot be reliably computed for arbitrary sequences. We therefore replaced the structural $\Delta\Delta G$ estimates with stability scores predicted by the transformer-based TemBERTure model [Rodella et al., 2024], which provides thermodynamic stability predictions directly from sequence. Each sequence in our dataset is thus annotated with two quantitative properties: (i) fluorescence brightness ($y_{\text{GFP}}$), experimentally measured by fluorescence-activated cell sorting, and (ii) thermodynamic stability ($y_{\text{stab}}$), predicted by TemBERTure. This representation enables consistent evaluation across both natural and generated sequences while maintaining comparability with the MosPro benchmark.

**Filtering Protocol.** To ensure methodological alignment with MosPro, we followed the same filtering logic applied in their fluorescence–stability benchmark, with the only modification being the use of TemBERTure stability scores instead of FoldX energies. Specifically, we first removed duplicate sequences and retained only variants with valid annotations for both fluorescence and stability. Let $Q_{0.4}(y_{\text{GFP}})$ and $Q_{0.4}(y_{\text{stab}})$ denote the 40th percentiles of fluorescence and stability, respectively. We then selected sequences satisfying

$$y_{\text{GFP}} \leq Q_{0.4}(y_{\text{GFP}}) \quad \text{and} \quad y_{\text{stab}} \leq Q_{0.4}(y_{\text{stab}}),$$

corresponding to the bottom 40% of both distributions. This selection follows the MosPro benchmark definition to ensure direct comparability while also defining a challenging low-fitness regime where simultaneous improvement of fluorescence and stability is nontrivial. Focusing on the lower 40% of both properties prevents trivial interpolation among already optimized sequences and encourages the model to explore upward optimization trajectories across the fitness landscape. Moreover, this regime yields a more balanced distribution of property values, reducing the natural skew toward bright and highly stable variants and reflecting realistic protein design scenarios in which both traits must be jointly optimized. Applying this filtering protocol yielded a final dataset of 9,769 sequences, which we used for subsequent embedding analysis, clustering, and structured sequence-pair generation.

**Embedding, Clustering, and Validation Split.** To characterize the diversity of protein sequences and construct a balanced dataset for model training, we first computed ESM2 embeddings [Lin et al., 2023] for all filtered sequences. ESM2 provides high-dimensional representations that encode evolutionary and biochemical relationships, enabling a meaningful comparison of sequence similarity in a learned functional space. Because the 1280-dimensional embeddings are difficult to interpret directly, we applied t-SNE to visualize their global structure in two dimensions.

The t-SNE projection (Fig. S1) reveals that sequences form multiple compact and well-separated regions, indicating distinct embedding patterns that likely correspond to different

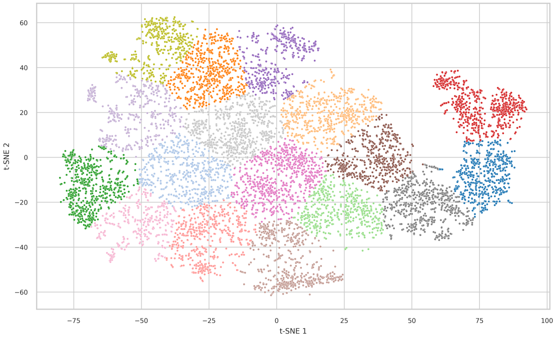


Fig. S1: Visualization of the t-SNE projection of ESM2 embeddings colored by the 17 $k$-means clusters. Each cluster captures sequences with similar embedding patterns, supporting balanced sampling across the latent space.

sequence families or structural motifs. To delineate these regions quantitatively, we applied $k$-means clustering to the t-SNE coordinates. Clustering serves two purposes: (1) to group sequences with similar embeddings for systematic sampling, and (2) to support structured pair generation for model training. By drawing pairs both within and across clusters, we can control the diversity of sequence pairs, which encourages the model to learn relationships that span both local and global regions of the embedding manifold. This structured pairing strategy reduces sampling bias toward dense areas and improves the representational coverage of the training data.

We set the number of clusters to $k = 17$ based on the observed structure of the t-SNE map. This value produced well-separated and approximately balanced clusters that captured the major modes of variation without overfragmenting similar regions. From each cluster, five sequences were randomly selected as the validation set, and the remaining sequences were used for training. This procedure ensures that the validation set spans the diversity of the embedding space and that each cluster contributes sequences for both training and validation. This clustering framework thus defines a structured organization of the embedding space that we leverage to generate training pairs with varying degrees of similarity. Specifically, we construct *in-cluster*, *cross-cluster*, and *random* sequence pairs to jointly capture local consistency and global diversity in the training data.

**Cross-Cluster Pair Generation.** We generated cross-cluster sequence pairs to provide broad property coverage and robust supervision under the following constraints:

1. **Cross-cluster sampling.** For each source sequence $i$ belonging to cluster $c$, one partner $j$ was sampled from each of the other clusters $c' \neq c$, producing up to sixteen candidate pairs per source.
2. **Uniqueness.** Each unordered pair is unique: if $(i, j)$ is created, the reverse pair $(j, i)$ is excluded.
3. **Adaptive thresholding.** To avoid degenerate pairs with negligible property differences, we introduced adaptive property-specific thresholds $\varepsilon_{y_{\mathrm{GFP}}}$ and $\varepsilon_{y_{\mathrm{stab}}}$, defined as
$$\varepsilon_x = 0.01 \times (\mathrm{P}_{99}(x) - \mathrm{P}_1(x)),$$
where $x \in \{y_{\mathrm{GFP}}, y_{\mathrm{stab}}\}$. Pairs were retained only if
$$|y_{\mathrm{GFP},i} - y_{\mathrm{GFP},j}| > \varepsilon_{y_{\mathrm{GFP}}} \quad \text{and} \quad |y_{\mathrm{stab},i} - y_{\mathrm{stab},j}| > \varepsilon_{y_{\mathrm{stab}}}.$$
This adaptive rule scales $\varepsilon$ with the empirical dynamic range of each property and effectively filters out trivial differences.
4. **Independent splits.** The same pairing procedure is applied independently to the training and validation partitions.

**In-Cluster Pair Generation.** To complement the cross-cluster sampling, we further constructed an *in-cluster* pairing dataset using the same adaptive-thresholding and filtering protocol described above. The only difference lies in the sampling strategy: for each source sequence $i$ in cluster $c$, we randomly sampled up to sixteen partners $j$ from within the same cluster ($c' = c$), excluding self-pairs and duplicates. All other settings including the adaptive thresholds, uniqueness constraints, and independent train/validation splits were kept identical to the cross-cluster procedure. This random pairs provide an unbiased control for evaluating whether the clustering-based sampling contributes to improved property coverage or optimization stability.

**Random Pair Generation.** In addition to the clustered sampling strategies, we also constructed a random pairing dataset to serve as a clustering-agnostic baseline. This procedure follows exactly the same adaptive-thresholding, filtering, and uniqueness settings as described above, but samples partners uniformly at random from the entire dataset without considering cluster assignments.

Figure S2 illustrates the distribution of absolute GFP score differences before applying the adaptive threshold, showing a strong concentration near zero. After applying the adaptive threshold, the distribution becomes more uniform, confirming that the thresholding strategy effectively removes near-identical pairs while maintaining sufficient diversity for learning.

We generated paired training data under multiple pairing strategies, each following the same adaptive-thresholding and filtering protocol described above. The cross-cluster and in-cluster procedures differ only in the sampling domain, across versus within clusters, while all other settings (uniqueness, adaptive thresholds, and independent splits) remain identical. In total, we obtained approximately 150k training pairs and 1k validation pairs for each scheme, as summarized in Table S2.

**Table S2.** Summary of generated pair datasets for the GFP–stability benchmark. Each pairing scheme uses the same adaptive thresholding and filtering protocol; only the sampling domain differs.

| Pairing scheme | Training pairs | Validation pairs |
|---|---|---|
| Cross-cluster | 154,944 | 1,360 |
| In-cluster | 152,224 | 1,267 |
| Random | 154,944 | 1,360 |

To assess how different pairing strategies affect optimization dynamics, we trained models independently on the cross-cluster, in-cluster, and random pair datasets described above. Figure S3 shows the evolution of the evaluation loss across global steps for each scheme. Among the three, the *in-cluster* pairing strategy achieves the lowest and most stable validation loss, while the *cross-cluster* scheme converges more slowly and exhibits higher fluctuations. The superior performance of the in-cluster pairs can be attributed to the smoother supervision signal arising from embedding-local relationships: pairs drawn within the same cluster tend to share similar sequence composition and local structure, resulting in more consistent property gradients and easier mapping

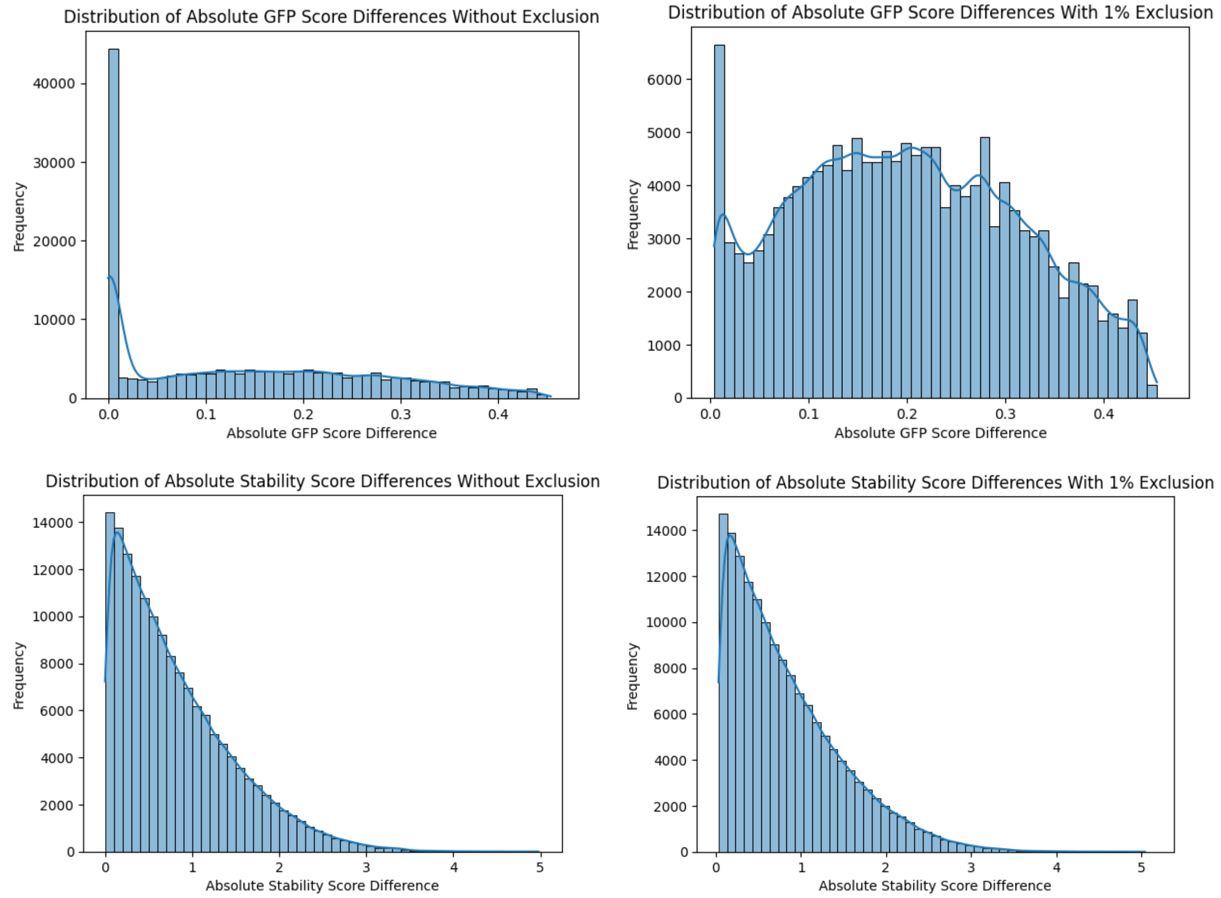


Fig. S2: Distributions of absolute property differences between paired sequences. (Top:) GFP brightness ($y_{\mathrm{GFP}}$) before (left) and after (right) applying the adaptive threshold. (Bottom:) Stability ($y_{\mathrm{stab}}$) before (left) and after (right) applying the adaptive threshold. For both properties, the adaptive thresholds (1% of the empirical range between the 99th and 1st percentiles) remove near-identical pairs, resulting in more uniform and informative distributions of pairwise differences.

between sequence differences and property changes. By contrast, cross-cluster pairs connect sequences that are distant in the embedding space and may differ across multiple structural or functional motifs, which introduces greater heterogeneity in the underlying property relationships. This diversity leads to noisier supervision and less stable optimization, as the model must reconcile large, often non-linear variations in both sequence and property space. The random pairing baseline lies between these two regimes, effectively mixing both local and global samples; it benefits from broader coverage but still inherits moderate variance from cross-cluster pairs. Overall, these observations suggest that the locality of pair sampling in the embedding space strongly influences learning stability and generalization, with in-cluster sampling providing the most coherent supervision signal.

#### F.2. Base Model: LoRA Fine-Tuned ProLLaMA

The ProLLaMA-7B model [Lv et al., 2025] is adopted as both the base model and the prior of the reward model. ProLLaMA is a protein language model trained in an autoregressive manner across diverse structural and functional protein superfamilies, capturing context-conditioned residue dependencies and long-range sequence patterns. This makes it suitable for protein design, where accurate modeling of sequence-function relationships and the underlying structural or biophysical constraints is essential.

To enable the ProLLaMA model to generate sequences belonging to the green fluorescent protein (GFP) superfamily, we fine-tuned the ProLLaMA base model using low-rank adaptation (LoRA). The LoRA approach enables parameter-efficient fine-tuning by injecting trainable low-rank matrices into specific layers of the pre-trained transformer while keeping the original model weights frozen. This significantly reduces GPU memory requirements while preserving model generalization capabilities.

**Dataset preparation.** The benchmark containing 51,715 unique GFP mutant sequences without property label (to prevent data leakage) was used for fine-tuning. We use "GFP Beta-barrel fluorescent protein superfamily" for this new superfamily to add, since it is close to an existing superfamily "beta-barrel domain superfamily" description. The sequences were tokenized using the same vocabulary and byte-pair encoding (BPE) tokenizer distributed with ProLLaMA, ensuring compatibility between the base model and the fine-tuned variant. The data were randomly split into training and validation sets in a ratio of 9:1 to monitor convergence and prevent overfitting.

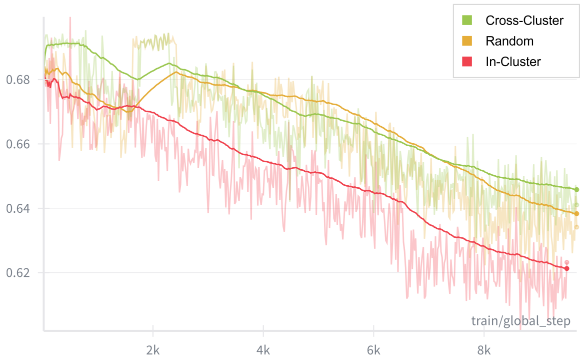


Fig. S3: Evaluation losses for different pairing strategies. The faint lines represent raw per-step validation losses recorded throughout training, and the solid curves show the exponentially smoothed averages for visualization. Each color corresponds to one pairing scheme: cross-cluster (green), random (yellow), and in-cluster (red). All models were trained under identical configurations, and the evaluation losses were computed on the same validation set. The smoothed curves highlight the general convergence trends and relative stability of each sampling strategy.

**Construction of instruction-following samples.** To align with the instruction-based training paradigm of ProLLaMA, each GFP sequence was converted into an instruction–response pair. Specifically, we designed synthetic text prompts that explicitly instructed the model to generate a sequence conditioned on a superfamily description. Each sample followed the textual template:

```
[Generate by superfamily]
Superfamily=<GFP Beta-barrel fluorescent protein
superfamily>
Answer:
Seq=<protein sequence>
```

This format enables the model to interpret the task as an instruction-following generation problem, consistent with its pretraining framework. Including the natural-language context improves conditioning on the intended superfamily and encourages generalization beyond direct sequence memorization, allowing the model to produce novel but structurally consistent GFP-like variants.

**Fine-tuning configuration.** Fine-tuning was performed using the `peft` and `transformers` libraries from Hugging Face, on 16 NVIDIA A100 GPUs (40 GB memory per GPU) with distributed data parallelism (DDP). The training employed the AdamW optimizer with a cosine learning rate scheduler. We used a base learning rate of $5 \times 10^{-5}$, effective batch size of 32 with gradient accumulation, and a LoRA rank of $r = 32$ with an $\alpha$ scaling factor of 64. The fine-tuning spanned approximately 2,800 update steps, corresponding to 1 epoch over the dataset. Mixed-precision training was employed, leveraging the `bfloat16` numerical format to improve computational efficiency and reduce GPU memory usage.

During fine-tuning, LoRA adapters were applied to the query and value projection matrices (`q_proj` and `v_proj`) of each transformer block in the model. The remaining parameters of the base ProLLaMA model were kept frozen throughout training.

**Model merging and storage.** After convergence, the learned LoRA adapter weights were merged ("baked") into the frozen base model weights to form a standalone model variant termed ProLLaMA-GFP-merged. The merged model thus integrates the GFP-specific representations while maintaining the general structural and functional knowledge of the original ProLLaMA. The merged model was finally uploaded to a private Hugging Face repository for reproducibility and long-term storage.

**Generation protocol.** To generate de novo GFP-like protein sequences, prompts were structured as the same like the format when used for training. Sampling used nucleus sampling ($p = 0.9$) with temperature $T = 0.8$ and a repetition penalty of 1.1. The model successfully produced diverse and coherent amino acid sequences consistent with the GFP structural motif.

**Comparison of distributions between generated and natural sequences.** To evaluate whether the model captured the residue-level characteristics of the GFP superfamily, we compared the positional amino acid distributions between sequences generated by GFP-finetuned ProLLaMA and the natural GFP sequences used during fine-tuning. A total of 5,000 sequences were sampled from the model using the generation protocol described above, and all natural sequences from the fine-tuning dataset were used for comparison.

All sequences were aligned by length and cropped to a common region corresponding to the canonical GFP beta-barrel domain. For each position, amino acid frequencies were computed and visualized using Logomaker [Tareen and Kinney, 2020], a Python library for creating sequence logos based on custom position–weight matrices. The total stack height at each position represents the information content derived from residue frequencies, while the relative letter heights correspond to the probability of each amino acid. This visualization provides an interpretable summary of positional conservation and sequence variability within the GFP superfamily.

Figure S4 displays the sequence logos for (a) the natural GFP sequences and (b) the GFP finetuned ProLLaMA generated sequences. The two logos exhibit highly similar residue distributions across most positions, with conserved motifs clearly recapitulated by the model. In particular, residues forming the beta-barrel scaffold and chromophore-binding pocket show similar amino acid preferences between the two sets. Minor deviations were observed in some regions, likely reflecting the model's generative variability rather than systematic drift.

These results demonstrate that the GFP finetuned ProLLaMA model successfully learned the sequence-level grammar of the GFP superfamily, enabling it to produce novel yet biochemically consistent variants that preserve key structural determinants of the native protein family.

In addition to the unconditioned generation analysis, we evaluated sequence-level properties of GFP variants generated under the trade-off preference-conditioned ST-PARM framework. All generated sequences were aligned to the wild-type GFP sequence used in the dataset, enabling direct position-wise comparison across different preference configurations. The resulting logos summarize residue frequency patterns at each position for preference-conditioned generations relative to the wild-type reference, revealing that preference conditioning induces localized and position-specific shifts in amino acid usage while

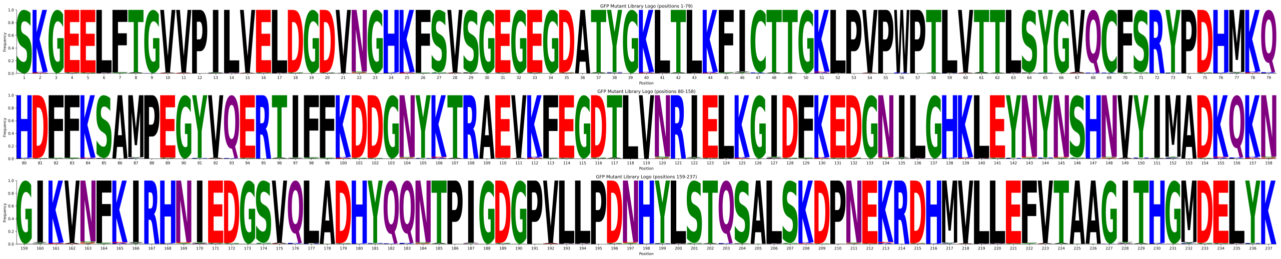

(a) Natural GFP sequences.

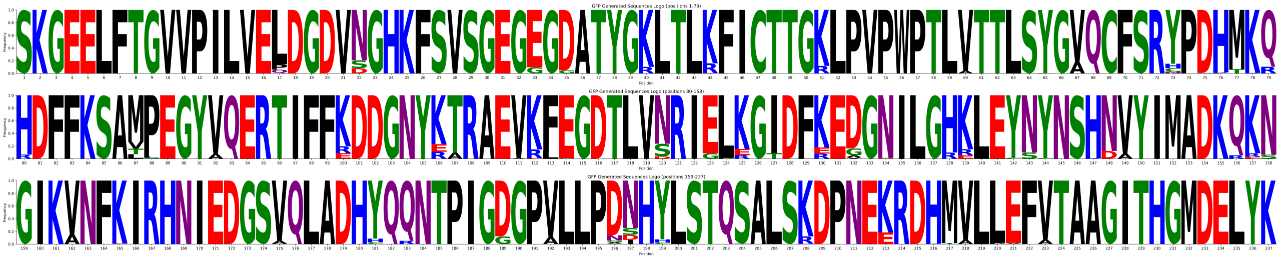

(b) ProLLaMA-GFP–generated sequences (unconditioned).

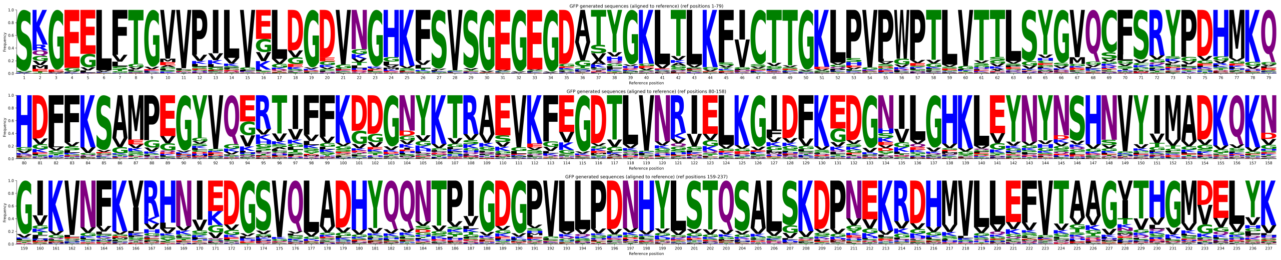

(c) ST-PARM–generated GFP sequences under different preference settings, aligned to the wild-type GFP reference.

Fig. S4: Sequence logo visualizations of amino acid distributions for GFP sequences. Logos display residue frequencies at each aligned position for natural sequences, unconditioned model generations, and preference-conditioned ST-PARM generations. All sequences are aligned to a common GFP reference where applicable.

**Table S3.** Oracle models used for property evaluation in the GFP design task. Each oracle provides sequence-level predictions that guide multi-objective optimization during ST-PARM training.

| Property | Oracle Model | Description |
|---|---|---|
| Fluorescence | GGS [Kirjner et al., 2024] | Energy-based model trained using contrastive divergence and Gibbs sampling. Assign each sequence a denoised fitness value from the learned energy function and apply graph-based smoothing to the empirical fitness landscape. |
| Thermodynamic Stability | TemBERTure [Rodella et al., 2024] | Transformer-based model that derives stability-relevant features from pretrained protein sequence embeddings. Maps these embeddings through a small prediction head trained on large-scale $T_m$ data to yield thermal stability estimates. |

conserved positions associated with the canonical GFP fold remain largely stable across preferences.

#### F.3. Oracle Models for Property Evaluation.

Two sequence-based oracle models were used to estimate the fluorescence and thermodynamic stability of designed GFP variants. Their key characteristics are summarized in Table S3.

#### F.4. Additional Results

Figure S5 reports the mean fluorescence and stability of generated sequences across preference settings, together with the PARM and MosPro baselines for reference.

Figure S6 reports the scatter plots of ST-PARM designs pre and post structural filtering.

Figure S7 shows that the retained Pareto frontiers under TM-score filter 0.7 and 0.8 shows similarity to that under TM score filter 0.5 in the main text.

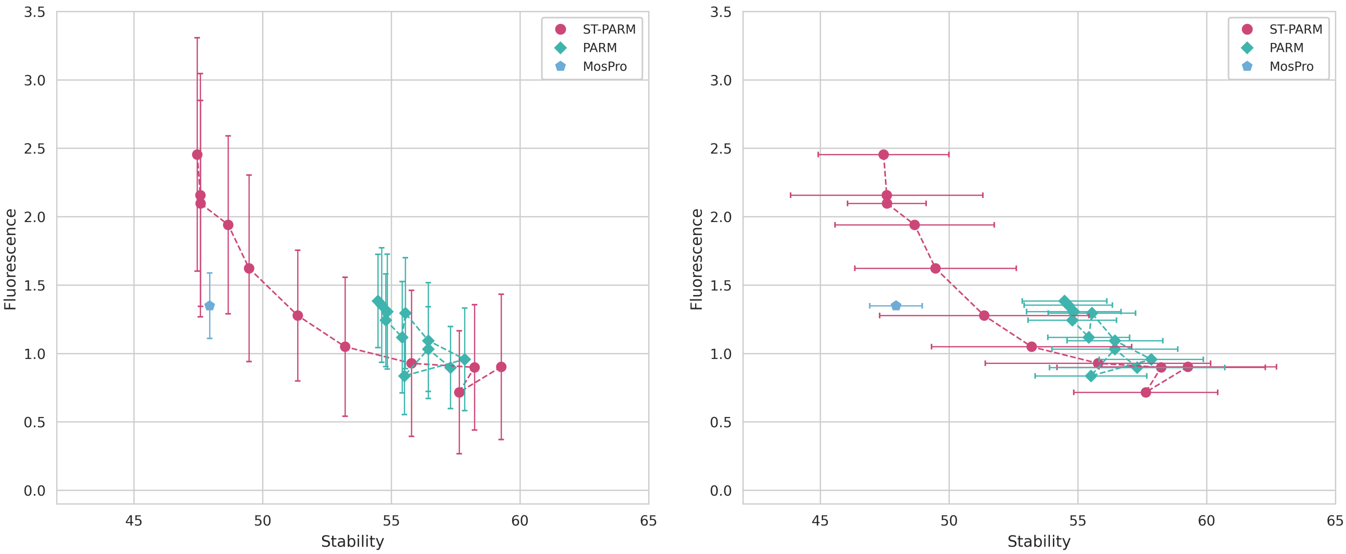


Fig. S5: Performance of ST-PARM on the GFP fluorescence–stability benchmark. Left: mean $\pm$ standard deviation of fluorescence for sequences generated under different trade-off preference vectors $\boldsymbol{\alpha} = (\alpha_{\text{fluor}}, \alpha_{\text{stab}})$. Right: corresponding stability scores under the same conditions. ST-PARM exhibits smooth and controllable transitions across preferences while achieving higher fluorescence and stability than the MosPro baseline. Error bars denote variation across independent generations.

##### F.5. Ablation Study: Reward Model Only Generation

**Motivation.** ST-PARM targets the aligned distribution

$$p_{\text{target}}(x \mid \alpha) \propto p_{\text{base}}(x)\, \exp(\beta R_\alpha(x)),$$

where the base LM provides an evolutionary prior and $R_\alpha$ reweights sequences toward Pareto-optimal regions. To assess the role of the prior, we perform an ablation in which generation is driven solely by the reward model.

**Setup.** Given a trade-off preference vector $\alpha$, sequences are optimized using reward-guided search without sampling from the LM. This removes all inductive biases learned from natural sequence distributions.

**Results.** Across the GFP stability–fluorescence task, removing the LM prior leads to:

- Irregular and unstable Pareto trade-offs. Reward-only frontiers vary erratically with $\alpha$ and lack the smooth progression produced by ST-PARM.
- Mode collapse toward extreme-stability regions. Sequences cluster tightly in high-stability regions, reflecting over-optimization of a single reward direction.
- Reward hacking and out-of-distribution drift. Designs achieve high predicted reward but leave the manifold explored by ST-PARM and the training data, indicating extrapolation errors in the reward model.
- Loss of evolutionary plausibility. Without $p_{\text{base}}$, the generator exploits weaknesses in $R_\alpha$ and produces unrealistic and repeating sequence patterns.

**Conclusion.** The reward model alone is not a generative mechanism. The base LM is essential for maintaining biological plausibility, stabilizing preference conditioning, and producing smooth, well-behaved Pareto frontiers.

#### G. IL-6 Nanobody Design: Protocols and Supplementary Results

##### G.1. Training Dataset Generation for Two-Objective Design

**Embedding and clustering for pairing strategy.** To construct informative training pairs, we first computed sequence embeddings for all IL-6 nanobody variants using the ESM2-650M protein language model [Lin et al., 2023]. These embeddings capture both structural and evolutionary relationships among sequences. We then applied UMAP (Uniform Manifold Approximation and Projection) to reduce the embeddings to two dimensions for visualization and neighborhood-based clustering. K-means Clustering was performed independently on the training and validation splits. Figure S9 shows the UMAP projections and clustering of ESM2 embeddings for the IL-6 nanobody sequences.

**Across-cluster vs. within-cluster pairing.** Based on the clustering results, we constructed two distinct pairing strategies: (i) within-cluster pairs, pairing sequences from the same cluster to emphasize local fine-grained variations, and (ii) across-cluster pairs, pairing sequences from different clusters to encourage global diversity and cross-structural generalization.

We evaluated models trained under each strategy on the evaluation set for across-cluster pairing. Figure S10 reports the evaluation loss trajectories for both configurations. The model trained on across-cluster pairs achieved consistently lower evaluation loss and faster convergence, suggesting that exposure to globally diverse sequence–property relationships improves generalization across structural manifolds. In contrast, within-cluster training led to overfitting to local sequence neighborhoods and higher evaluation loss on unseen cross-cluster comparisons.

##### G.2. Oracle Models for Two-Objective Evaluation.

Two sequence-based oracle models were employed to estimate the biophysical properties of candidate nanobody sequences: one

**Table S4.** Oracle models used for property evaluation in the IL-6 nanobody design task. Each oracle provides sequence-level predictions that guide multi-objective optimization during ST-PARM training.

| **Property** | **Oracle Model** | **Description** |
|---|---|---|
| Thermodynamic Stability | DeepSTABp [Jung et al., 2023] | Deep neural network trained on large-scale protein datasets to predict protein thermal stability (melting temperature, $T_m$) directly from amino acid sequences. Captures sequence–stability relationships through convolutional and transformer layers. |
| Solubility | CamSol [Sormanni et al., 2015] | Physics- and statistics-informed sequence model estimating intrinsic solubility propensity from residue hydrophobicity, charge, and sequence context. Produces normalized solubility scores inversely related to aggregation tendency. |

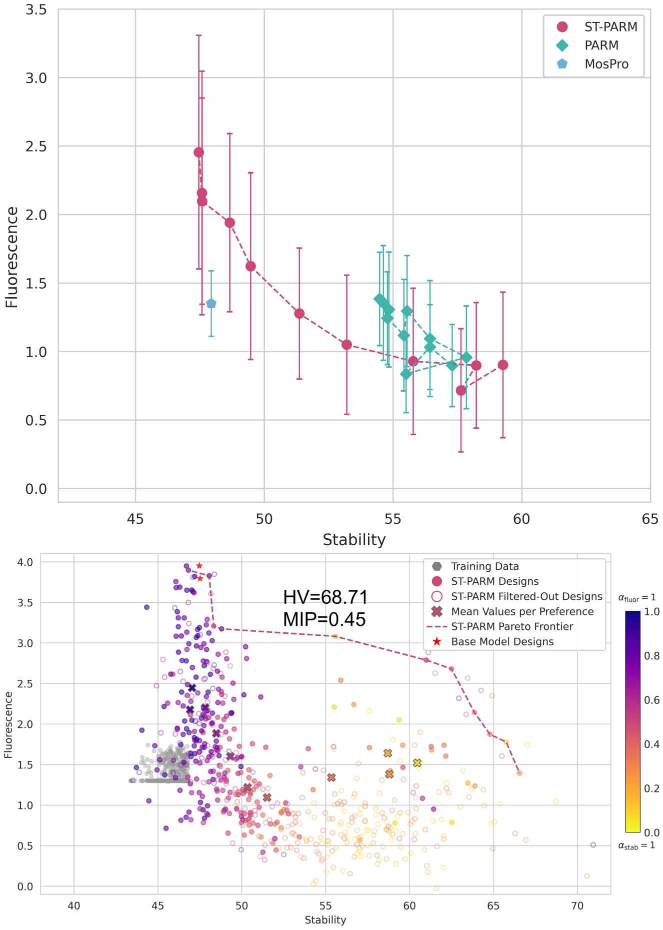


Fig. S6: ST-PARM designs (pre and post structural filtering) on the GFP fluorescence–stability benchmark. Structural filtering (local confidence and global fold preservation) yields a conservative, interpretable filter that complements fluorescence–stability objectives, while maintaining high Pareto coverage (post vs. pre HV=68.71 vs. 74.65) and tradeoff controllability (post vs. pre MIP=0.45 vs. 0.44).

for thermodynamic stability and one for solubility. Their key characteristics are summarized in Table S4.

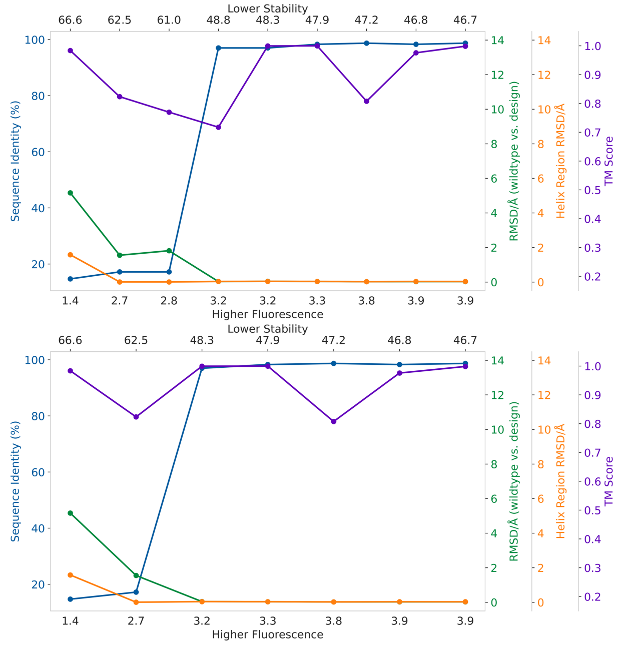


Fig. S7: Structural variation along the GFP fluorescence–stability Pareto frontier (filtered with pLDDT⩾80 and TM scores ⩾0.7 (top) or 0.8 (bottom). Reported are sequence identity, TM scores, backbone RMSD (TM-aligned region), and chromophore-containing helix RMSD of representative frontier designs relative to wild-type avGFP, as the preference to fluorescence increases toward the right.

#### G.3. Three-Objective Design: Stability–Solubility–Affinity

To further assess the generality of ST-PARM for higher-dimensional design problems, we extend the IL-6 nanobody task to a three-objective optimization setting involving predicted stability, solubility, and binding affinity. This experiment complements the two-objective study presented in the main text and demonstrates the capability of preference-conditioned inference to navigate three-way trade-offs among developability and functional constraints.

**Training data construction.** The original dataset includes a binary binding label for each nanobody. To exploit this information for pairwise preference learning, we construct preference pairs by pairing every binding sequence with every non-binding sequence.

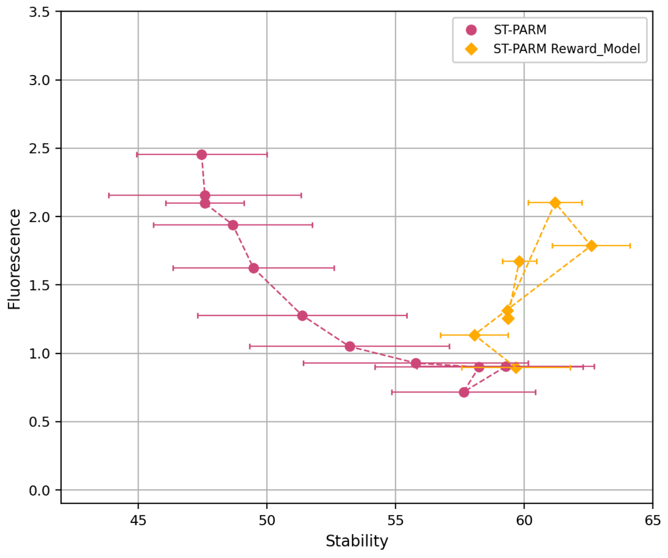


**(A)** Preference sweep with stability std.

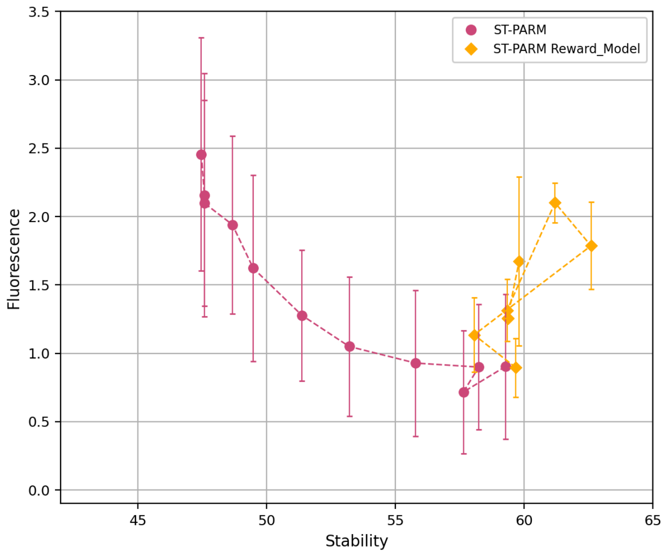


**(B)** Preference sweep with fluorescence std.

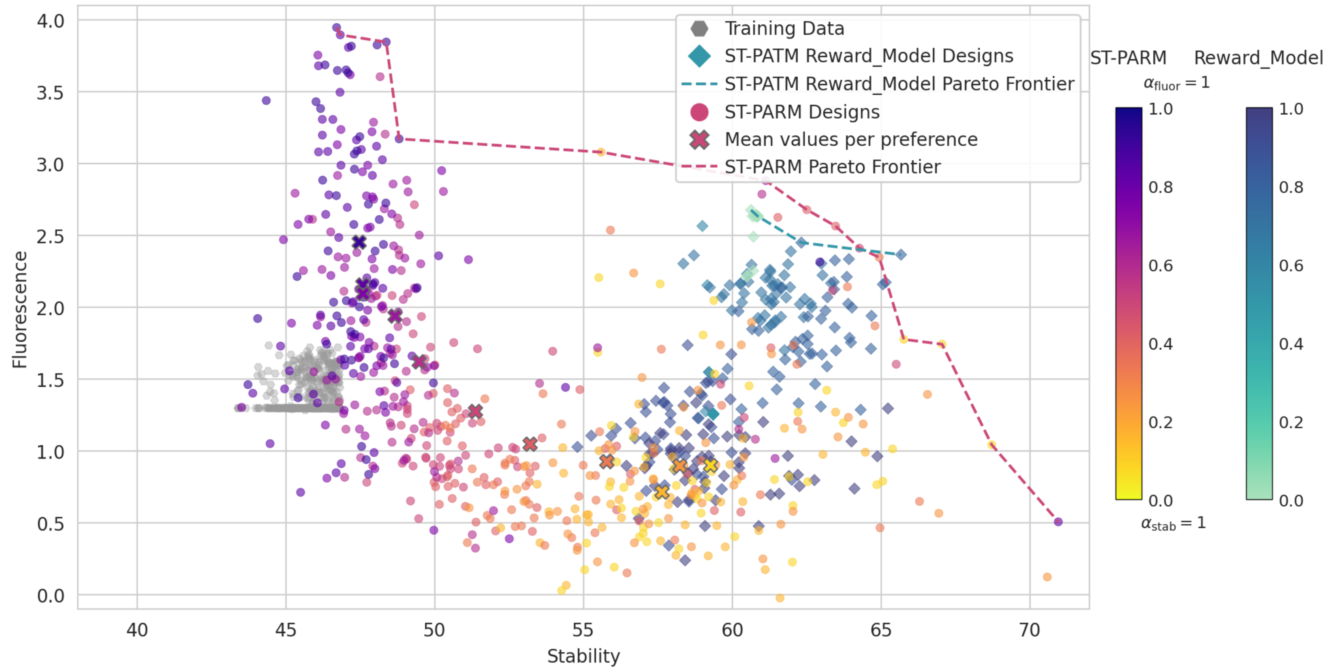


**(C)** Scatter plot comparing stability–fluorescence trade-offs for ST-PARM versus reward-model-only generation.

Fig. S8: Comparison of ST-PARM versus reward-model-only generation. Removing the LM prior produces unstable trade-offs, collapsed diversity, and reward-hacking behavior.

**Objective-specific oracle models.** Each of the three objectives in the IL-6 nanobody task is evaluated using a dedicated sequence-based oracle model. Thermostability is predicted using TEMPRO [Alvarez and Dean, 2024], a protein thermostability estimator using language model embeddings. Aggregation propensity (solubility) is evaluated with TANGO [Rousseau et al., 2006], which quantifies the intrinsic tendency of a sequence to form aggregation-prone motifs. Binding affinity is assessed using the BindPred model [Piao et al., 2025], a sequence based binding affinity predictor. For each designed nanobody, BindPred scores its affinity against all 31 IL-6 antigen variants, and the affinity objective is taken as the maximum predicted value across these antigens.

**Three-objective evaluation.** For a sweep of trade-off preference vectors $\alpha \in \Delta^2$, ST-PARM generates nanobody candidates that reflect different trade-offs among the three objectives. We visualize the resulting Pareto-structured design space using three pairwise projections, each colored by the third objective:

1. stability–solubility plane, colored by affinity;
2. stability–affinity plane, colored by solubility;

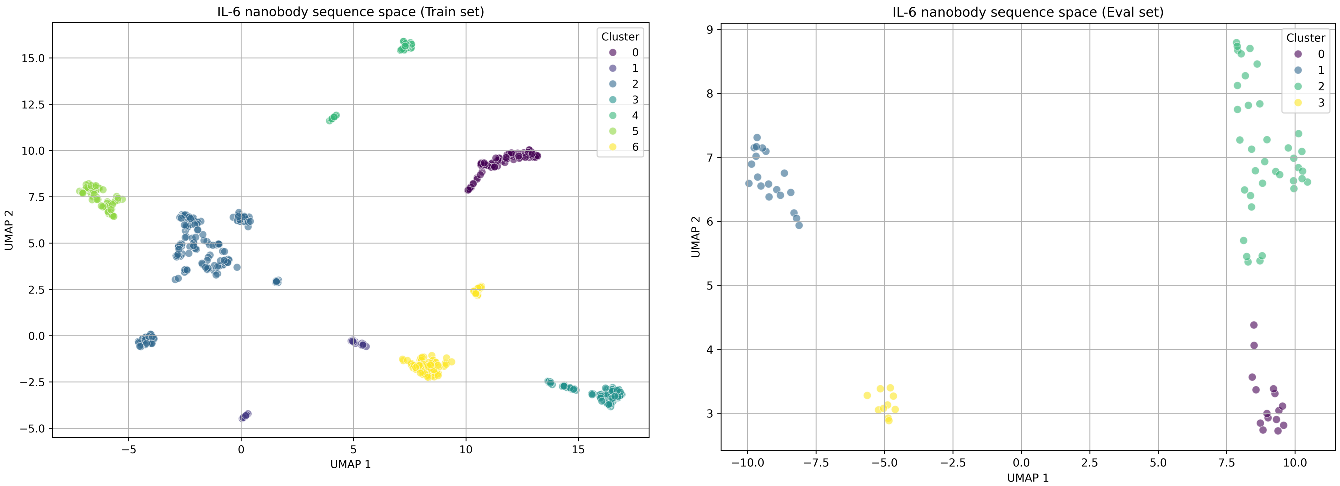


**(A)** Training set clustering.

**(B)** Evaluation set clustering.

Fig. S9: UMAP projections of ESM2 embeddings for IL-6 nanobody sequences. Two-dimensional projections were obtained using UMAP on ESM2 embeddings, followed by kNN-based clustering applied separately to the training (A) and evaluation (B) splits. Colors denote cluster membership.

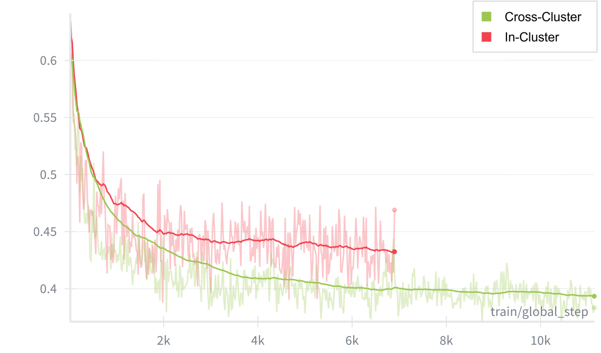


Fig. S10: Evaluation losses on the across-cluster test set. Models trained with across-cluster pairs (green) exhibit lower losses and better generalization compared to those trained with within-cluster pairs (red), indicating that diverse pairing improves robustness across structural clusters.

3. solubility–affinity plane, colored by stability.

These complementary views reveal how ST-PARM traverses and interpolates across three competing objectives while maintaining smooth preference responsiveness. Representative results are shown in Figure S11.

##### G.4. Evaluator Robustness Results for Three-Objective Designs

We assess the robustness of the designs to the choice of objective evaluators as in Figure S12. The positive correlations across evaluator choices support the robustness.

#### H. Safety–Helpfulness Alignment on PKU-SafeRLHF

For generality, we also assess ST-PARM on the PKU-SafeRLHF-10K benchmark, a two-objective alignment task in natural language generation. Each prompt is paired with completions annotated for *safeness* and *helpfulness* with each of objective evaluator, representing competing behavioral goals in instruction tuning.

##### H.1. Dataset and Training

We adopt the PKU-SafeRLHF-10K dataset [Ji et al., 2024] for evaluating multi-objective alignment in natural language generation. This dataset consists of 10,000 human-annotated prompt–response pairs designed for safety alignment of large language models. Each sample includes a user query $x$ and two candidate responses $(y_1, y_2)$. We augmented it with two continuous labels indicating how each response is scored under two separate criteria: *helpfulness* and *harmlessness* (or safeness); We use two LM models as the scorers. The dual-objective structure enables simultaneous optimization of informativeness and behavioral safety, providing a controlled benchmark. Following [Lin et al., 2025a], we split the dataset into 8,000 samples for training, 500 for validation, and 1,500 for testing, as summarized in Table S5.

We train a preference-conditioned autoregressive reward model on annotated tuples $(x, y_1, y_2)$, where $x$ denotes a user prompt and $(y_1, y_2)$ are paired responses labeled for *safeness* and *helpfulness*. The model is optimized using multi-objective supervision, which is the smooth Tchebycheff scalarization of the per-objective (reward-calibrated) losses. The LLaMA-2-7B model [Touvron et al., 2023] serves as both the frozen base model and the initialization backbone of the reward model. Larger language models can similarly be used as the base model, incurring only additional inference cost. At inference, responses are generated across a sweep of trade-off preference vectors $(\alpha_{\text{help}}, \alpha_{\text{safe}}) \in \Delta^1$.

##### H.2. Oracle Models for Objective Evaluation

Two separate reward (oracle) models were used to provide scalar signals for the objectives of helpfulness and safeness. Both models are part of the PKU-Alignment `Beaver-7B-v1.0` suite [Dai

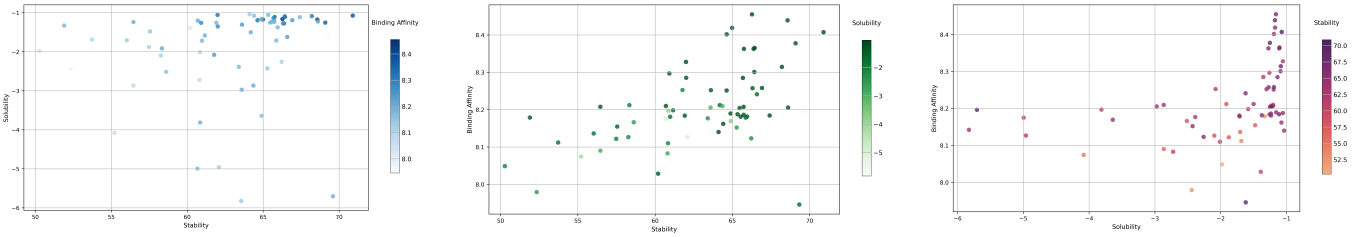


**(A)** Stability–solubility projection (colored by affinity).

**(B)** Stability–affinity projection (colored by solubility).

**(C)** Solubility–affinity projection (colored by stability).

Fig. S11: Three-objective IL-6 nanobody design visualized across pairwise objective projections.

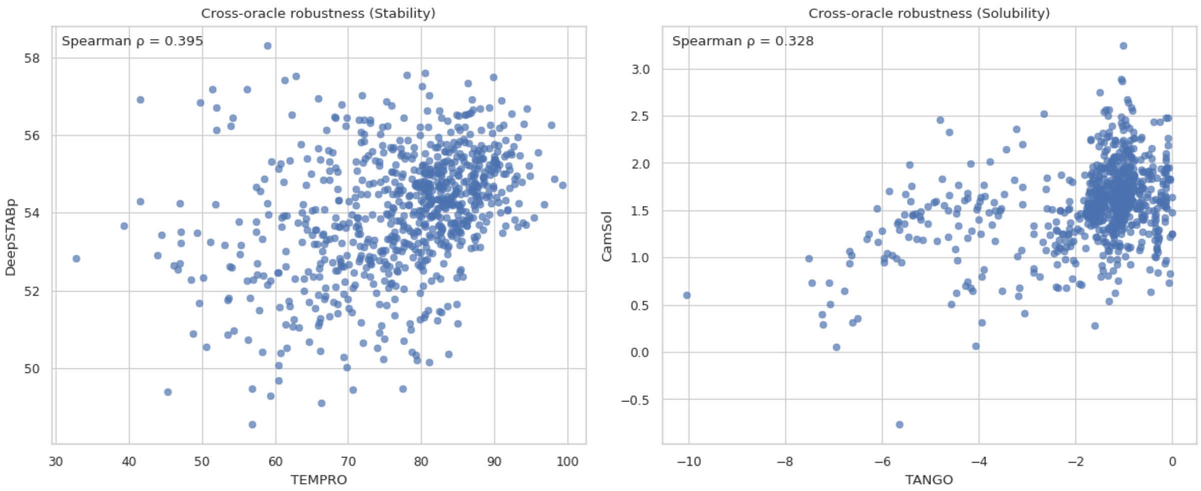


Fig. S12: Evaluating designs with alternative evaluators (oracles) shows positive rank-correlations between the two choices for either stability or solubility, supporting the robustness to evaluator choice.

et al., 2023], fine-tuned on human preference annotations that disentangle safe behavior from informative and relevant responses.

#### H.3. Results

We study how ST-PARM's performance depends on (i) the training prior distribution over trade-off preference vectors $\alpha$ and (ii) the smoothing parameter $\tau$ in the smooth Tchebycheff scalarization, then compare against baselines. Across all analyses, the best outcomes arise from *intermediate* settings; neither overly concentrated nor overly diffuse $\alpha$ priors, and neither vanishingly small nor overly large $\tau$. It reflects a balance between coverage of the Pareto frontier and stable gradient signals.

**Effect of the trade-off preference prior.** According to the smooth Tchebycheff scalarized loss in Eq. S12, we train a series of ST-PARM models under different training priors for the trade-off preference vector $\alpha = (\alpha_{\text{help}}, \alpha_{\text{safe}})$, where each component is sampled from a symmetric Beta($\texttt{param}, \texttt{param}$) distribution with varying concentration parameter `param`. This setup controls how frequently the model encounters extreme versus balanced trade-offs during optimization. Figure S13(a) illustrates the induced marginal distributions over $\alpha_{\text{help}}$ (or equivalently $\alpha_{\text{safe}}$), while Figure S13(b) shows the corresponding gradient-norm statistics measured during training. Figure S14 further reports the preference-swept averages of per-objective scores at a fixed smoothing parameter $\tau = 0.5$, and table and figure S15 summarizes the resulting Hypervolume (HV) and Mean Inner Product (MIP). Together, these results reveal how the choice of $\alpha$ distribution affects both the optimization dynamics and the final multi-objective performance.

When the concentration parameter `param` is too small, although it is well optimized for the extreme preference regions so the span of the Pareto curve is large, it produces sparse coverage and poor generalization in the balanced trade-off regions; This can be showed in the Figure S14: compared to the more balanced cases (warm color curves), the density of the points corresponding to the interior, balanced regions of the preference simplex (when inference-time $\alpha$ is near (0.5, 0.5)) is lower for these more extreme cases (cold color curves).

One hypothesis is that, during training in these small `param` cases, the sampled preferences on the 1-simplex $\Delta^1$ are dominated

**Table S5.** Statistics of the PKU-SafeRLHF-10K dataset. The dataset provides pairs of responses to the prompt.

| **Dataset** | **Train pairing #** | **Validation pairing #** | **Test pairing #** |
|---|---|---|---|
| PKU-SafeRLHF-10K | 8,000 | 500 | 1,500 |

**Table S6.** Oracle models used for the Safety–Helpfulness alignment task. Each model provides scalar label estimates corresponding to a distinct alignment objective.

| **Objective** | **Oracle Model (Path)** | **Description** |
|---|---|---|
| Helpfulness | PKU-Alignment/beaver-7b-v1.0-reward | An evaluation model fine-tuned from LLaMA-7B on human preference data emphasizing informativeness, relevance, and instruction-following quality. Produces scalar helpfulness scores aligned with human judgments. |
| Safeness | PKU-Alignment/beaver-7b-v1.0-cost | A companion cost model trained to predict unsafe or harmful behavior likelihoods. Lower scores indicate safer responses; higher scores penalize toxicity, bias, or policy violations. Used as the safeness oracle by negating the cost signal. |

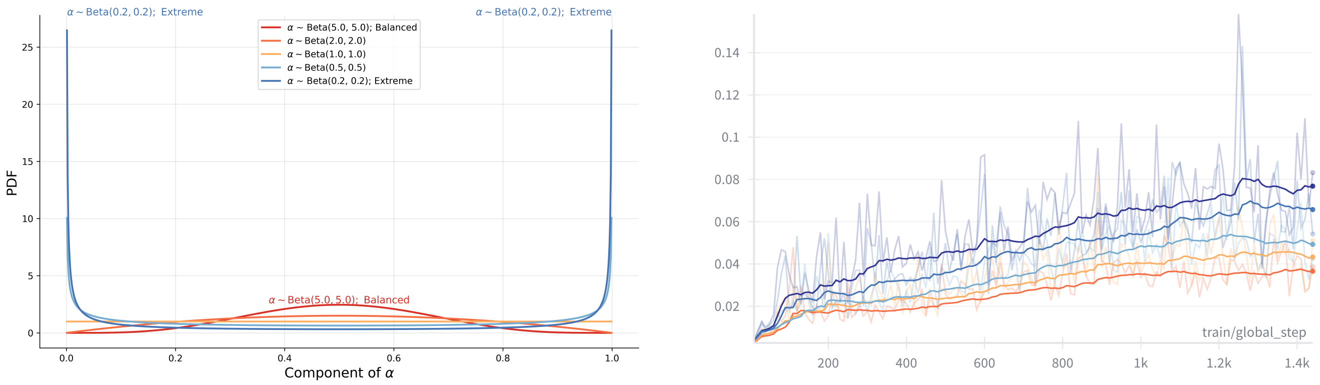


**(A)** PDFs of $\alpha$ components under Beta($\texttt{param}, \texttt{param}$). Small $\texttt{param}$ values bias toward extreme preferences, large values cluster near balanced trade-offs, and intermediate values balance both.

**(B)** Gradient-norm statistics during training under different preference priors. Intermediate Beta$(0.5, 0.5)$ maintains informative yet stable gradients, while extreme or overly concentrated priors lead to unstable or vanishing updates.

Fig. S13: Effect of preference prior on optimization dynamics. (A) Distributions of sampled preference weights $\alpha$. (B) Corresponding gradient-norm behavior during ST-PARM training. Intermediate concentration parameters yield balanced gradient magnitudes and stable convergence.

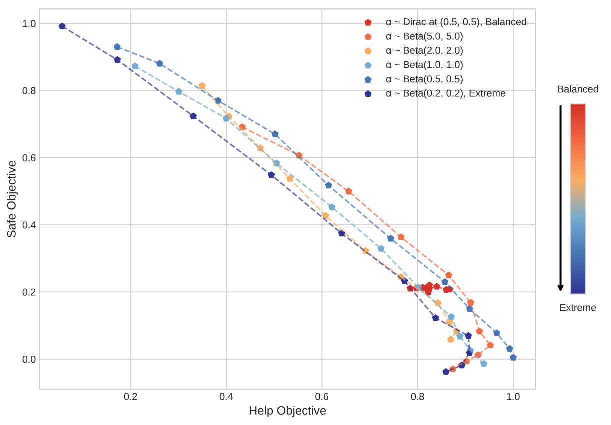


Fig. S14: Effect of the $\alpha$ prior on averaged objective scores. We fix $\tau$ at 0.5. Intermediate $\texttt{param}$ yields smoother preference tracking and better front coverage than overly extreme or overly central priors.

by extreme, near-corner samples, corresponding to almost single-objective optimization as we can see in Figure S13(a). Although this leads to rapid improvement on the extremes, such biased sampling causes the model to receive strong and unidirectional gradient signals however emphasizing one objective at a time. The fast changing and in large magnitude gradient in this case can be seen in the Figure S13(b). As a result, the model under-explores regions where gradients from different objectives must be balanced, limiting its ability to learn smooth trade-offs across the Pareto front.

Conversely, when the concentration parameter $\texttt{param}$ is large, the sampled preferences become highly concentrated around the simplex center, as shown in Figure S13(a). In this regime, gradients from the competing objectives often point in opposing directions and partially cancel, resulting in small overall gradient magnitudes and slow optimization progress, as confirmed in Figure S13(b). Consequently, the model primarily optimizes for the balanced regions of the preference space while neglecting the extremes, leading to limited exploration. As illustrated in Figure S14, this manifests as clustered trade-off trajectories (warm-colored curves), with reduced sensitivity to changes in $\alpha$ compared to models trained under more extreme priors (cold-colored curves). In the limiting case of a Dirac distribution at $\alpha = (0.5, 0.5)$, the learned responses collapse, yielding highly clustered and degraded performance that effectively flattens the model's response to varying preferences.

*Summary.* An *intermediate* concentration parameter $\texttt{param}$ provides the most effective compromise: it exposes the model to

| Trade-off preference $\alpha$ Distribution | HV | MIP |
|---|---|---|
| Dirac (0.5, 0.5) | 22.19 | 0.51 |
| Beta(5.0, 5.0) | 58.13 | 0.64 |
| Beta(2.0, 2.0) | 57.51 | 0.66 |
| Beta(1.0, 1.0) | 58.46 | 0.69 |
| Beta(0.5, 0.5) | **64.78** | **0.74** |
| Beta(0.2, 0.2) | 53.87 | 0.69 |

**(A)** Numerical results. Hypervolume (HV) and Mean Inner Product (MIP) for models trained with different $\alpha$ priors.

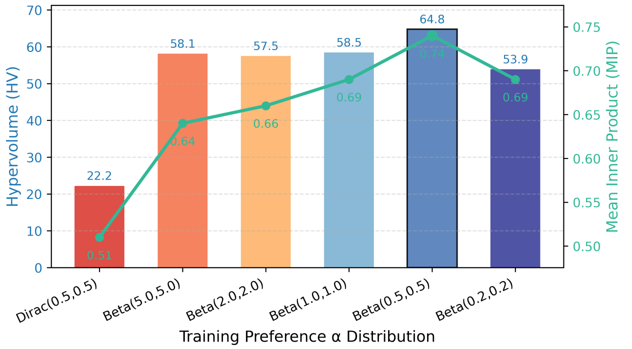


**(B)** Visual summary. HV and MIP as a function of the Beta(`param`, `param`) concentration parameter.

Fig. S15: Effect of training preference distribution on ST-PARM performance.

both extreme and balanced trade-offs, maintaining informative gradient magnitudes throughout training and resulting in higher hypervolume (HV), and stronger preference alignment (MIP).

**Effect of the smoothing parameter $\tau$.** We next study how the smooth Tchebycheff (STCH) smoothing parameter $\tau$ governs optimization dynamics and inference-time controllability. Recall that the STCH loss takes the form:

$$\mathcal{L}_{\mathrm{STCH}}(\alpha) = \tau \log \sum_{i=1}^{k} \exp\Big(\tfrac{\alpha_i(\ell_i - z_i)}{\tau}\Big) \tag{S17}$$

whose gradient with respect to the per-objective losses can be written as a softmax-weighted convex combination:

$$\frac{\partial \mathcal{L}_{\mathrm{STCH}}}{\partial \ell_i} = \underbrace{\frac{\exp\big(\frac{\alpha_i(\ell_i - z_i)}{\tau}\big)}{\sum_j \exp\big(\frac{\alpha_j(\ell_j - z_j)}{\tau}\big)}}_{w_i(\alpha,\ell,\tau)} \cdot \alpha_i.$$

As shown in Figure S16, when $\tau \to 0$, $\mathcal{L}_{\mathrm{STCH}}$ approaches the hard Tchebycheff operator and $w_i$ concentrates on the maximal (scaled) deviation, recovering a Pareto-complete operator that aggressively targets the current worst objective. Conversely, as $\tau \to \infty$, the $\mathcal{L}_{\mathrm{STCH}}$ approaches a weighted sum, dampening contrast among objectives and smoothing curvature.

*Small $\tau$.* When $\tau$ is too small, $w_i(\alpha, \ell, \tau)$ becomes nearly one-hot on the dominant objective for the current mini-batch. This yields large gradient norms and sharp updates (Figure S16), rapidly improving corner solutions but providing limited signal in interior regions. Empirically, the $\alpha$-sweep shows pronounced movement at preference extremes with reduced sensitivity in balanced trade-offs (Figure S17), and the resulting Pareto coverage underrepresents non-extreme segments.

*Large $\tau$.* When $\tau$ is too large, $w_i$ becomes nearly uniform (modulated by $\alpha$ only weakly through the numerator), which suppresses contrast between objectives. Optimization remains numerically stable (small gradients), but learning resembles linear scalarization and in theory tends to blur non-convex regions of the frontier, diminishing responsiveness to changes in $\alpha$. This manifests as smoother yet compressed trade-off trajectories (Figure S17) and reduced HV, along with plateaued MIP due to weak trade-off preference tracking.

*Intermediate $\tau$.* At intermediate values (e.g., $\tau \approx 0.5$ in our setting), $w_i$ preserves sufficient contrast to guide progress toward the current most underperforming objective while retaining enough smoothing to stabilize gradients. This balance produces (i) moderate, informative gradient magnitudes throughout training (Figure S16); (ii) smooth inference-time responses across the $\alpha$-sweep (Figure S17); and (iii) improved HV and MIP (Figure S18), indicating broader Pareto coverage and stronger preference adherence.

*Summary.* The two limits of $\tau$ expose opposing failure modes: very small $\tau$ concentrates updates on the single worst objective, causing unstable learning; very large $\tau$ flattens objective contrast and degrades sensitivity to preference variation, acting like linear scalarization. An intermediate $\tau$ preserves enough selectivity to navigate non-convex front regions while ensuring stable gradients, yielding the best HV/MIP trade-off and the most faithful preference controllability.

**Comparison with baselines.** Figure S19 reports the mean *helpfulness* and *safeness* achieved across a sweep of inference-time preference weights $\alpha \in \{(0.0, 1.0), (0.1, 0.9), \ldots, (1.0, 0.0)\}$ for PARM [Lin et al., 2025a], MOD [Shi et al., 2024], ST-PARM, and its reward-calibrated variant. Under the intermediate configuration (`param` $= 0.5$, $\tau = 0.5$) identified above, ST-PARM exhibits smoother and more coherent trade-off trajectories, leading to denser coverage of the Pareto frontier compared to other methods. The reward-calibrated variant maintains comparable overall behavior while achieving a further increase in frontier coverage. Quantitative results in Table S7 corroborate these trends: averaged over the full preference sweep, ST-PARM substantially improves both the hypervolume (HV) and mean inner product (MIP) relative to PARM and MOD, and the reward-calibrated variant attains the highest aggregate scores across both metrics.

*Mechanistic interpretation.* The superior performance of ST-PARM arises from its use of a smooth Tchebycheff scalarization, which provides a differentiable relaxation of the max-based Pareto objective. The additional reward calibration reweights each pairwise loss according to the observed label difference, ensuring that larger label gaps contribute stronger learning signals and reducing noise from ambiguous comparisons.

*Comparison with PARM.* Both ST-PARM and PARM train a single preference-conditioned autoregressive reward model (ARM) that jointly encodes all objectives. In PARM, preference weights are incorporated via a linear scalarization and parameter-efficient PBLoRA adaptation, providing compact storage and low inference

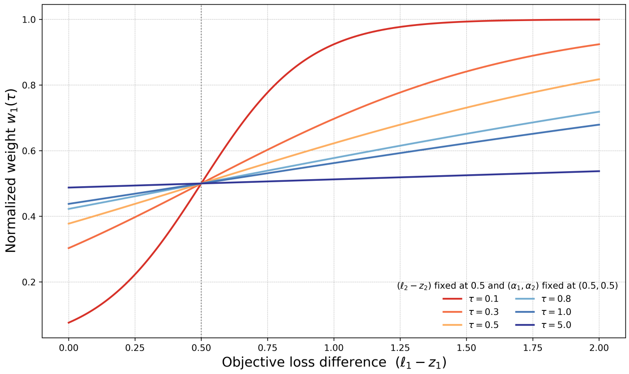


**(A)** Induced per-objective weighting $w_1(\tau)$ under different smoothing parameters. Smaller $\tau$ yields sharply peaked weights (hard selection of the dominant loss), while larger $\tau$ flattens the weighting toward uniform averaging. Intermediate $\tau$ maintains a relatively smooth but contrastive focus, balancing stability and sensitivity.

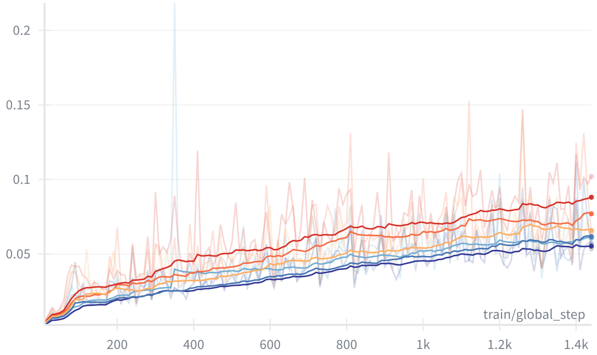


**(B)** Training-time gradient norms under different smoothing parameters $\tau$. Small $\tau$ (sharp scalarization) leads to large, volatile gradients dominated by a single objective, while large $\tau$ (over-smoothed) produces small, less informative gradients. An intermediate $\tau$ yields stable yet responsive optimization dynamics.

Fig. S16: Effect of smoothing parameter $\tau$ on optimization behavior. (A) The induced weighting distribution $w_1(\tau)$ shows how $\tau$ controls the sharpness of objective prioritization in the smooth Tchebycheff operator. (B) Corresponding gradient norms during training highlight the trade-off between instability (small $\tau$) and over-smoothing (large $\tau$).

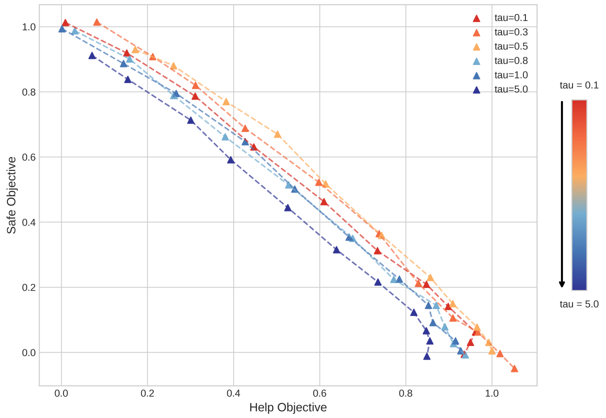


Fig. S17: Effect of $\tau$ on preference-swept average scores. Small $\tau$: unstable, corner-focused; large $\tau$: over-smoothed, under-sensitive. Intermediate $\tau$ maximizes controllability and coverage.

cost relative to training multiple independent ARMs. Unlike the linear scalarization in PARM, which enforces proportional trade-offs, the non-linear smooth Tchebycheff objective in ST-PARM dynamically accentuates dominant losses, yielding smoother interpolation and improved Pareto frontier.

*Comparison with MOD.* MOD adopts a decoding-based composition strategy, forming preference-conditioned responses by combining next-token probabilities from multiple single-objective models in proportion to the desired trade-off preference vector, derived analytically under an $f$-divergence–regularized objective. This approach enables flexible, training-free control over trade-offs but necessitates simultaneous access to multiple models, increasing memory requirements. Moreover, its theoretical guarantees apply at the sequence level, while the practical token-wise implementation can introduce local inconsistencies and hinder Pareto optimality. Empirically, these characteristics manifest as reduced HV compared to unified training approaches such as PARM and ST-PARM.

*Quantitative interpretation.* Across the complete $\alpha$ sweep (Table S7), ST-PARM raises HV from approximately 53.0 (for both PARM and MOD) to 64.78, with a corresponding increase in MIP to 0.74. Incorporating reward calibration further elevates HV to 73.65 and MIP to 0.78. These improvements substantiate that unified preference conditioning, when coupled with smooth Tchebycheff scalarization and calibrated reward scaling, leads to more complete Pareto coverage and stronger adherence to user-specified trade-offs. By contrast, MOD achieves competitive but less consistent interpolation due to its token-level approximation, while PARM delivers stable yet worse coverage limited by its linear scalarization.

**Table S7.** HV and MIP on PKU-SafeRLHF. Scores are averaged over the full $\alpha$ sweep.

| **Model** | **HV** | **MIP** |
|---|---|---|
| PARM [Lin et al., 2025a], | 53.39 | 0.72 |
| MOD [Shi et al., 2024] | 53.28 | 0.64 |
| ST-PARM (Ours) | 64.78 | 0.74 |
| ST-PARM + Reward-Calibrated (Ours) | **73.65** | **0.78** |

| **Smoothing parameter $\tau$** | **HV** | **MIP** |
|---|---|---|
| 0.1 | 58.38 | 0.73 |
| 0.3 | 62.29 | **0.75** |
| 0.5 | **64.78** | 0.74 |
| 0.8 | 54.93 | 0.71 |
| 1.0 | 54.87 | 0.71 |
| 5.0 | 49.18 | 0.66 |

**(A)** Numerical results. Hypervolume (HV) and Mean Inner Product (MIP) as a function of the smoothing parameter $\tau$.

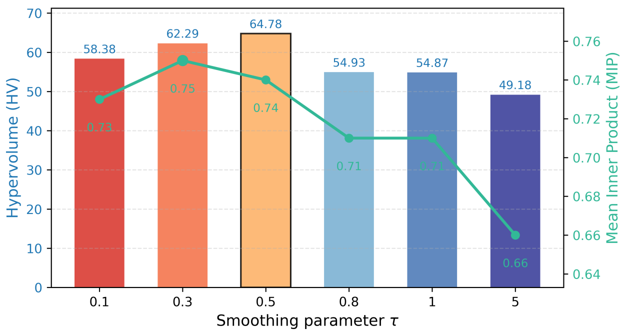


**(B)** Visual summary. Bars indicate HV and the line indicates MIP.

Fig. S18: Effect of the smoothing parameter $\tau$ on ST-PARM performance. (A) Numerical comparison of HV and MIP across different $\tau$ values. (B) Corresponding visual trends for a fixed preference prior $\alpha \sim \text{Beta}(0.5, 0.5)$. Intermediate $\tau$ achieves the best balance between gradient stability, Pareto coverage, and preference alignment.

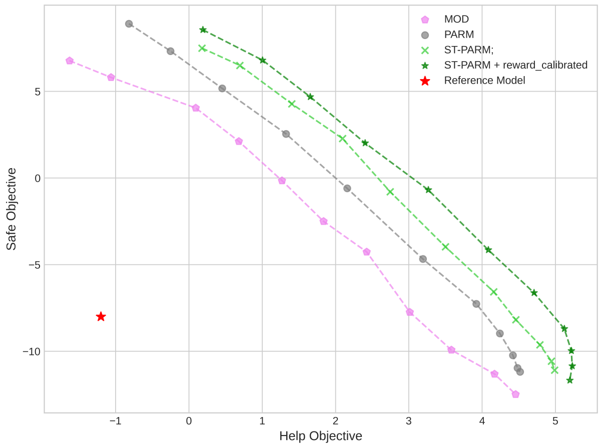


Fig. S19: Trade-off curves under varying inference-time preferences. ST-PARM (and its reward-calibrated variant) provide smoother interpolation and tighter trade-off preference tracking than PARM and MOD.
